# B/T cell crosstalk and aberrant inflammatory IgG exacerbate autoimmune intestinal inflammation

**DOI:** 10.1101/2022.09.12.507066

**Authors:** Iana Gadjalova, Julia M. Heinze, Marie Christine Goess, Julian Hofmann, Julian J. Albers, Ria Spallek, Birgit Blissenbach, Annalisa Buck, Marie-Christin Weber, Emely Scherer, Maximilian Kampick, Rupert Öllinger, Oleg Krut, Roland Rad, Katja Steiger, Christof Winter, Klaus-Peter Janssen, Philipp-Alexander Neumann, Raif S. Geha, Jürgen Ruland, Selina J. Keppler

## Abstract

Dysregulated B cell responses have been described in inflammatory-bowel disease (IBD) patients; however, the role of B cells in IBD pathology remained incompletely understood. We here described Wiskott-Aldrich Syndrome interacting protein deficient (*Wipf1*^*-/-*^) mice as novel mouse model of spontaneous, chronic colitis modelling human IBD. Concomitant with aberrant IgG production in colonic tissue of *Wipf1*^*-/-*^ mice, we identified systemic, hypo-sialylated IgG as drivers of IL-1β production in monocytes. Pathological antibody production was promoted by the hyper-reactivity of *Wipf1*^*-/-*^ B cells in response to LPS stimulation, resulting in efficient activation of the MAPK/Erk and mTOR/Akt/4E-BP1 pathways and heightened metabolic activity. In addition to abundant inflammatory IgG, we found that B cells directly promoted the production of pro-inflammatory cytokines by intestinal CD4^+^ T cells. B/T co-culture assays defined the co-stimulatory molecule CD86 as driver of IFN-γ and GM-CSF production by CD4^+^ T cells. CD86 expression was further enhanced by the presence of sCD40L, which was elevated in sera of *Wipf1*^*-/-*^ mice. Similarly, colonic B cells of IBD patients expressed increased mRNA levels of CD86 correlating with enhanced levels of systemic sCD40L. Together, B cell-mediated pro-inflammatory cytokine secretion and B cell-derived inflammatory antibody production contributed to exacerbated pathogenesis during intestinal inflammation.

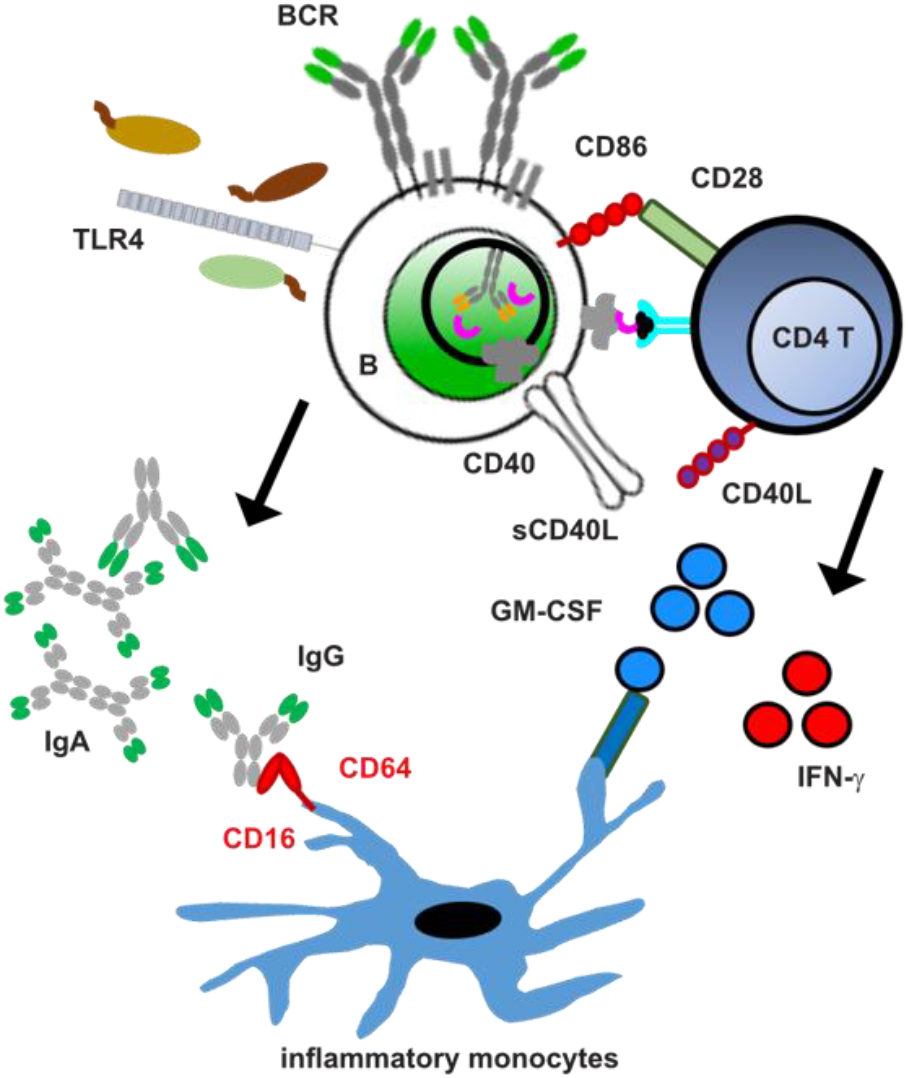

**One Sentence Summary:** B cells fuel intestinal inflammation

## INTRODUCTION

Inflammatory bowel disease (IBD), including Crohn’s disease and ulcerative colitis (UC), is a group of immune-mediated disorders of the intestine with increasing incidence worldwide (*1*). Susceptibility to IBD is driven by environmental risk factors together with a genetic predisposition to aberrant mucosal responses to commensals, including the production of pro-inflammatory cytokines by multiple T cell subsets, as well as macrophages (*2-4*). B cells might influence these responses through interaction with T cells or antibody production.

In the intestine, activation of B cells takes place in mesenteric lymph nodes (mLN), Peyer’s Patches (PP) and isolated lymphoid follicles (ILF) of the lamina propria (LP) and occurs via both T-independent (TI) and T-dependent (TD) pathways. TI responses potentially involve the recognition of microbial components prevalent in the gut such as lipopolysaccharide (LPS) through Toll-like receptor 4 (TLR4) expressed on B cells. TD responses require signals from CD4 T follicular helper (TfH) cells (*5*), which provide co-stimulatory signals for B cell differentiation and class-switch recombination (CSR) via ligands that include CD40L (*6*). During inflammation, exaggerated immune reactions to the microbiota involve a TD response, leading to antibodies of IgA as well as IgG isotypes (*7*). Recent scRNA sequencing studies revealed perturbations in humoral immunity during IBD, including the accumulation of B cells, an IBD-specific subset of TfH cells and IgG plasma cells (PC) in mucosal tissue (*8, 9*). IgG-antibodies produced against the microbiota correlated with disease severity and have been described to bind to Fc gamma receptors (FcγR)-expressing monocytes leading to Interleukin (IL)-1β secretion and sustained intestinal inflammation (*10*). However, the role of B cells and antibody-secreting cells (ASC) in IBD pathology remains elusive.

A large genome-wide association study among IBD patients identified over 163 loci associated with IBD risk (*11*). A network analysis including these risk loci as well as gene expression data identified an IBD sub-network that contains several genes - amongst them the gene encoding for the actin cytoskeleton regulator Wiskott-Aldrich Syndrome protein (WASp). WAS is an X-linked immunodeficiency associated with an increased susceptibility of infections and autoimmune manifestations. 10% of WAS patients develop early-onset IBD (*12*), accompanied by systemic autoimmunity, such as IgA nephropathy (*13*). Causes of disease are defective expression of the WASp or the WASp interacting protein (WIP), both regulators of the actin cytoskeleton. WASp as well as WIP-deficient (*Wipf1*^*-/-*^) mice develop IBD with 100% penetrance (*14, 15*). Murine *Wipf1*^*-/-*^ B cells are hyper-proliferative to both BCR and TLR signaling and demonstrate enhanced class-switch recombination to IgG1 after TLR4 stimulation *in vitro* (*16*). We thus hypothesized that the presence of hyper-reactive B cells in the gut might enhance inflammation during IBD development.

Our results suggest that, during autoimmune intestinal inflammation, hyper-reactive B cells in the gut provide critical co-stimulatory signals to CD4 T cells and differentiate into antibody-secreting cells which provide pro-inflammatory IgG. We here provide evidence for B cells as drivers of IBD pathogenesis and suggest that our mouse model can be of immense value to test treatment options targeting B cells during intestinal inflammation.

## RESULTS

### *Wipf1*^*-/-*^ mice model human chronic colonic inflammation

*Wipf1*^*-/-*^ mice develop signs of intestinal inflammation as early as 9 weeks of age, as apparent by a lengthening and thickening of the colon (Fig. S1A, Fig. S1C) as well as lymphadenopathy in colon draining mesenteric lymph nodes (colonic mLN, the first and last node of the mLN chain(*17*)) and enlarged spleens (Fig. S1B). Characterizing intestinal inflammation in *Wipf1*^*-/-*^ mice in detail, we found cell infiltration into the mucosa and submucosa, concomitant with the appearance of neutrophils and mast cells; a thickening of the muscularis interna as well as irregular crypts with increased crypt length, crypt abscesses and goblet cell loss, which resulted in a colitis score with mild to moderate severity (Fig. 1A+B, Fig. S1D, Table S1 (*18*)). In addition, we found an increased relative mRNA expression of the proinflammatory cytokines IL-6 and IL-17A in the inflamed colonic LP of *Wipf1*^*-/-*^ mice (Fig. S1E). To determine whether intestinal inflammation is associated with an altered composition of the commensal microbiota, 16S ribosomal RNA sequencing was performed on total bacteria isolated from cecal content. Strikingly, while younger (<9 weeks) *Wipf1*^*-/-*^ mice showed similar bacterial diversity compared to controls, we observed a reduced alpha diversity (Shannon index) in older (>11 weeks) *Wipf1*^*-/-*^ mice compared to age-matched controls (Fig. 1C). Younger *Wipf1*^*-/-*^ mice demonstrated similar microbial composition compared to WT controls (Jaccard distance), whereas older *Wipf1*^*-/-*^ mice differed significantly (Fig. S1F-I) and contained relatively less *Lachnospiracea* and relatively more *Lactobacillacea* (Fig. 1D, S1H+I). *Wipf1*^*-/-*^ mice hence demonstrate characteristic features of colonic inflammation with an onset of disease at 9 weeks and a manifestation at 11 weeks of age.

**Figure 1:**
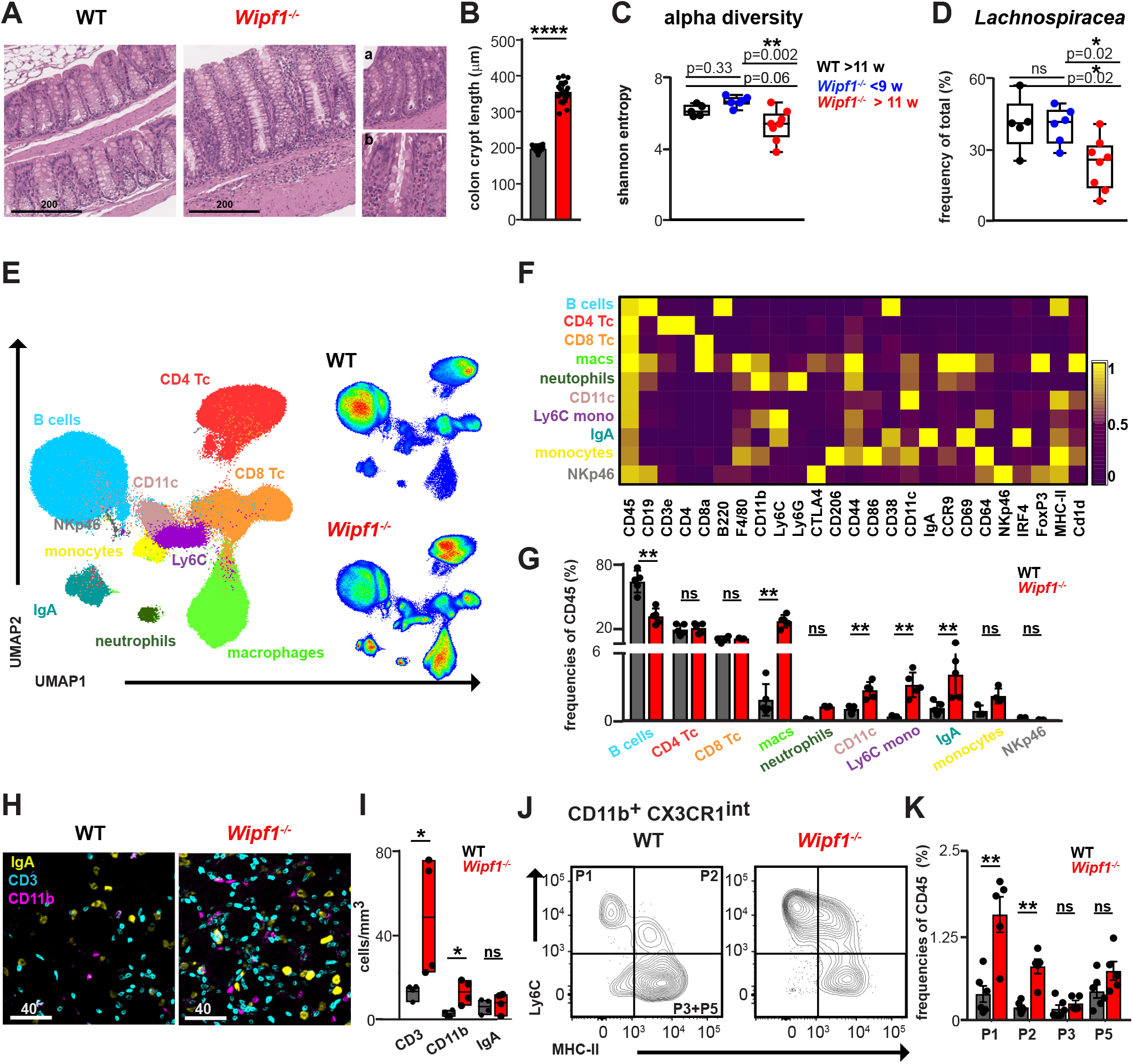
*Wipf1*^*-/-*^ mice model human chronic colonic inflammation. **(A)** Representative pictures of colonic sections of WT or *Wipf1*^*-/-*^ mice stained for H&E. Zoom-ins of sections of *Wipf1*^*-/-*^ mice demonstrate **a**. crypt abscesses and **b**. neutrophil infiltration (scale bar 200 μm). **(B)** Colonic crypt length measured from sections stained with H&E (total of 3 images, 20 crypts each, mean ± SD). Statistical significance was calculated using student’s t test. **(C+D)** Analysis of fecal bacteria from 11 week old WT and young (<9 weeks) or older (>11 weeks) *Wipf1*^*-/-*^ mice. (**C)** Fecal bacteria alpha diversity (Shannon entropy) is shown. (**D)** Abundances of *Lachnospiracea*. Representative data from one of two independent experiments giving similar results (Box and whisker blots (min to max)). Non-parametric ordinary one-way ANOVA with multiple comparisons was used to test for significant differences. **(E-G)** Lymphocytes isolated from colonic tissue were analyzed by mass cytometry. **(E)** Uniform Manifold Approximation and Projection for Dimension Reduction (UMAP) was used to depict immune cell populations in the CD45 expressing population. FlowSOM-based immune cell populations are overlaid as color dimension. **(F)** Normalized mean population expression levels of all markers used for UMAP visualization and FlowSOM clustering. Frequencies of **(G)** immune cell lineages of WT and *Wipf1*^*-/-*^ mice as defined by the FlowSOM clustering (n=5 per genotype, pooled from 2 independent experiments, mean ± SD). Statistical significance was calculated using Mann-Whitney U-test. **(H)** Immunofluorescent microscopy of cleared colonic tissue pieces of WT and *Wipf1*^*-/-*^ mice stained for IgA plasma cells, CD3 T cells and CD11b monocyte macrophages (scale bar 40 μm). **(I)** Quantification of immunofluorescent images (total of 4 mice each with 1 3D image each, floating bars (min to max), bar indicates mean). Statistical significance was calculated using student’s t test. **(J+K)** Flow cytometric analysis of colonic “monocyte waterfall” subsets isolated from WT or *Wipf1*^*-/-*^ mice. **(J)** Representative flow blots of waterfall subsets pre-gated on CD45+, CD19-, CD4-, CD11b+, CX3CR1int. **(K)** Frequencies of respective waterfall subsets (n=5 per genotype, pooled from 2 independent experiments, mean ± SEM). Statistical significance was calculated using Mann-Whitney U-test. *p < 0.05, **p < 0.01, ***p < 0.001, ****p < 0.0001, ns = not significant

To assess the recruitment and activation of immune cells to the site of inflammation, we isolated CD45 expressing lymphocytes from colonic tissue and analyzed these cells for the protein expression of several lineage- and activation associated surface markers as well as transcription factors using mass cytometry (Fig. 1E-G). We found an overabundant population of innate immune cells (monocytes/macrophages) as well as an increased population of IgA PC in lymphocytes isolated from the colonic LP (Fig. 1G) and mLN (Fig. S1J-L) of *Wipf1*^*-/-*^ compared to WT mice.

To gain insight into the distribution of lymphocytes in the intact tissue, we used cleared colonic tissue pieces together with volumetric imaging (Hofmann et al). We detected infiltration of CD3 T cells as well as CD11b monocyte/macrophage populations in colonic crypts of the mucosa of *Wipf1*^*-/-*^ compared to WT mice (Fig. 3H+I). Intestinal macrophages largely derive from Ly6C^hi^ monocytes infiltrating the LP that progressively lose their Ly6C but increase MHC-II expression (P1-P4 subsets, CD11b^+^CX3CR1^+^ ‘‘monocyte waterfall’’ (*19*). Using flow cytometry, we observed an influx of colonic Ly6C^hi^ MHC-lI^low^ (P1) monocytes and an increase in newly differentiated inflammatory Ly6C^+^MHC-II^int^ (P2) macrophages in the colonic tissue of *Wipf1*^*-/-*^ compared to WT mice (Fig. 1J+K, Fig. S3M-Q).

Together these findings show that *Wipf1*^*-/-*^ mice demonstrate characteristic features of colonic inflammation, concomitant with a reduced microbial diversity and enhanced accumulation of T cells and inflammatory monocytes/macrophages in colonic tissue.

### Aberrant colonic as well as systemic humoral B cell responses in *Wipf1*^*-/-*^ mice promote monocyte activation

Using immunohistochemistry as well as immunofluorescence of colonic swiss rolls, we observed large B cell-containing lymphoid follicles in the colon of *Wipf1*^*-/-*^ mice that were rarely detected in littermate controls. These follicles also contained T cells, indicative of an ongoing TD immune response (Fig. 2A, S2A+B). We found an accumulation of similar follicles in sections of inflamed colonic tissue of IBD patients compared to non-inflamed colonic tissue (Fig. S2C). While total cell numbers as well as frequencies of CD19 B cells were similar (Fig. S2D+E), we found a 3-fold increase in the frequency of germinal center (GC) B cells expressing GL-7 and CD95 in lymphocytes isolated from the *Wipf1*^*-/-*^ colonic LP compared to WT controls (Fig. 2B+C, S2F).

**Figure 2:**
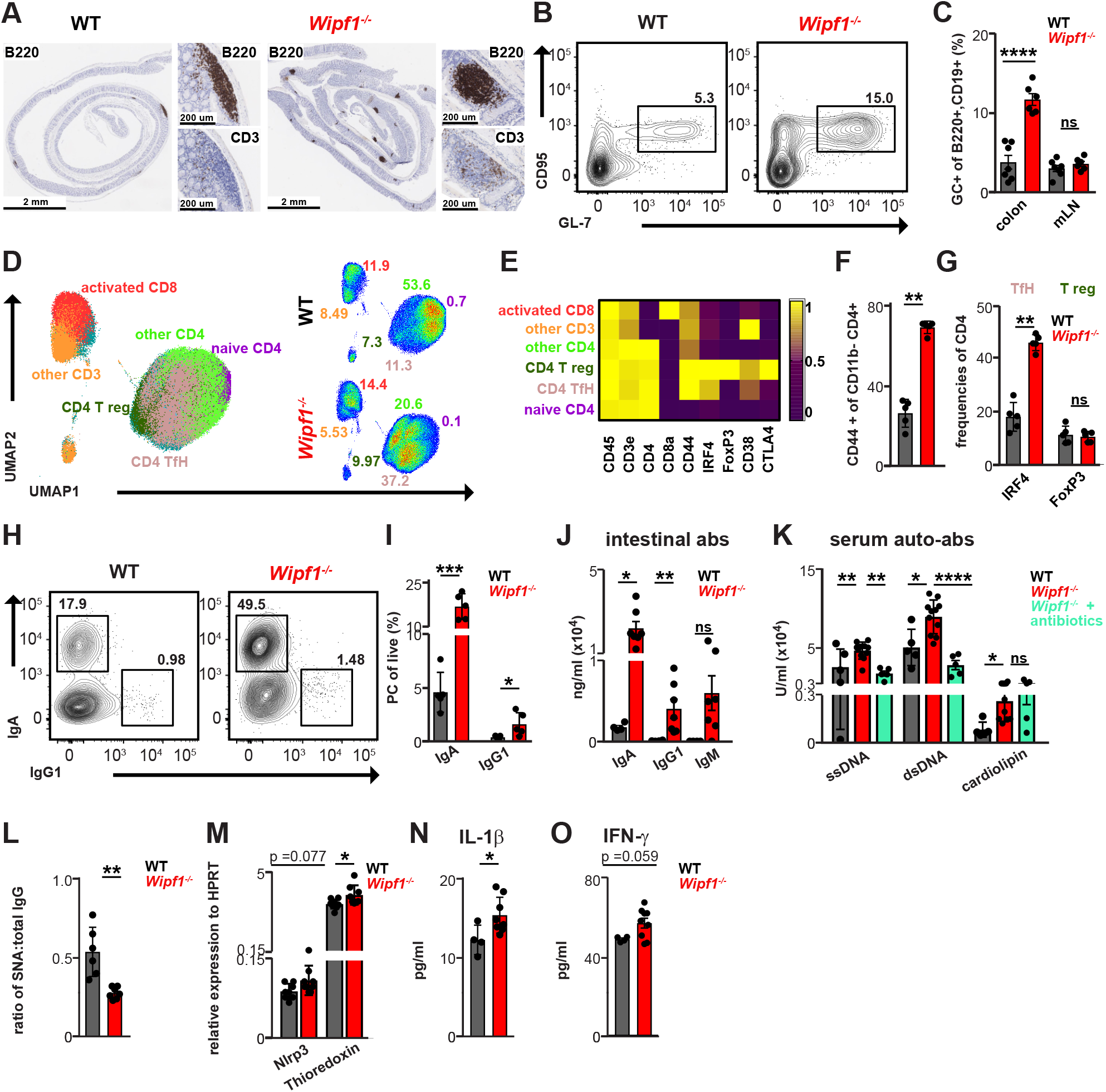
Aberrant colonic as well as systemic humoral B cell responses in *Wipf1*^*-/-*^ mice promote monocyte activation. **(A)** Immunohistochemistry of sections of colonic swiss rolls stained for B220. Zoom-ins of isolated lymphoid follicles (ILFs) indicating B220 as well as CD3 staining. **(B+C)** Lymphocytes isolated from colonic tissue of WT or *Wipf1*^*-/-*^ mice were stained to determine germinal center B cells (GC B cells; B220+GL7+CD95+). **(B)** Representative flow blots of GC B cells **(C)** Frequencies of GC B cells of total B cells from colonic tissue or mesenteric lymph nodes (mLN) (n≥6 per genotype, pooled from at least 3 independent experiments, mean ± SEM). Statistical significance was calculated using student’s t test. **(D-G)** Lymphocytes isolated from colonic tissue of WT or *Wipf1*^*-/-*^ mice were analysed by mass cytometry. **(D)** UMAP was used to depict CD45 and CD3 expressing T cell population. FlowSOM-based immune cell populations are overlaid as color dimension. **(E)** Mean population expression levels of T cell markers used for UMAP visualization and FlowSOM clustering. Frequencies of **(F)** activated CD4 T cells and **(G)** IRF4+ T follicular helper (TfH) or FoxP3+ regulatory T cells (Tregs) (n=5 per genotype, pooled from 2 independent experiments). Statistical significance was calculated using Mann-Whitney U-test. **(H+I)** Lymphocytes isolated from colonic tissue were stained to determine plasma cells (PC) using flow cytometry **(H)** Representative flow blots of IgG1 or IgA expressing PC **(I)** Frequencies of PC of total live cells from colonic tissue (n≥5 per genotype, pooled from 2 independent experiments, mean ± SEM) Statistical significance was calculated using Mann-Whitney U-test. **(J)** Indicated antibody isotypes were detected in mucus scrapes of WT or *Wipf1*^*-/-*^ mice using a multiplex assay **(K)** Indicated auto-antibodies detected in serum of WT or *Wipf1*^*-/-*^ mice or *Wipf1*^*-/-*^ mice treated with antibiotics using ELISA (n≥5 per genotype, mean ± SEM) Statistical significance was calculated using student’s t test. **(L)** Levels of (2,6)-sialic acid residues on total serum IgG were measured by ELISA. IgG sialylation is expressed as ratio of SNA binding (OD) to the concentration of total IgG (n≥6 per genotype, mean ± SD). Statistical significance was calculated using student’s t test. **(M)** Bone-marrow derived monocytes (BMDMs) were incubated with total purified serum IgG of WT or *Wipf1*^*-/-*^ mice for 20 hrs. Relative mRNA expression normalized to HPRT of *Nlrp3* and *thioredoxin* determined by qPCR **(N)** IL-1β production by WT BMDMs primed for 16 hrs with LPS followed by 4 hrs incubation with purified serum IgG and 30 min ATP stimulation. **(O)** IFN-Ψ production by WT BMDMs primed for 4 hrs with LPS and purified serum IgG **(L-O)** n≥5 sera per genotype, mean ± SEM. Statistical significance was calculated using student’s t test. *p < 0.05, **p < 0.01, ***p < 0.001, ****p < 0.0001, ns = not significant

**Figure 3:**
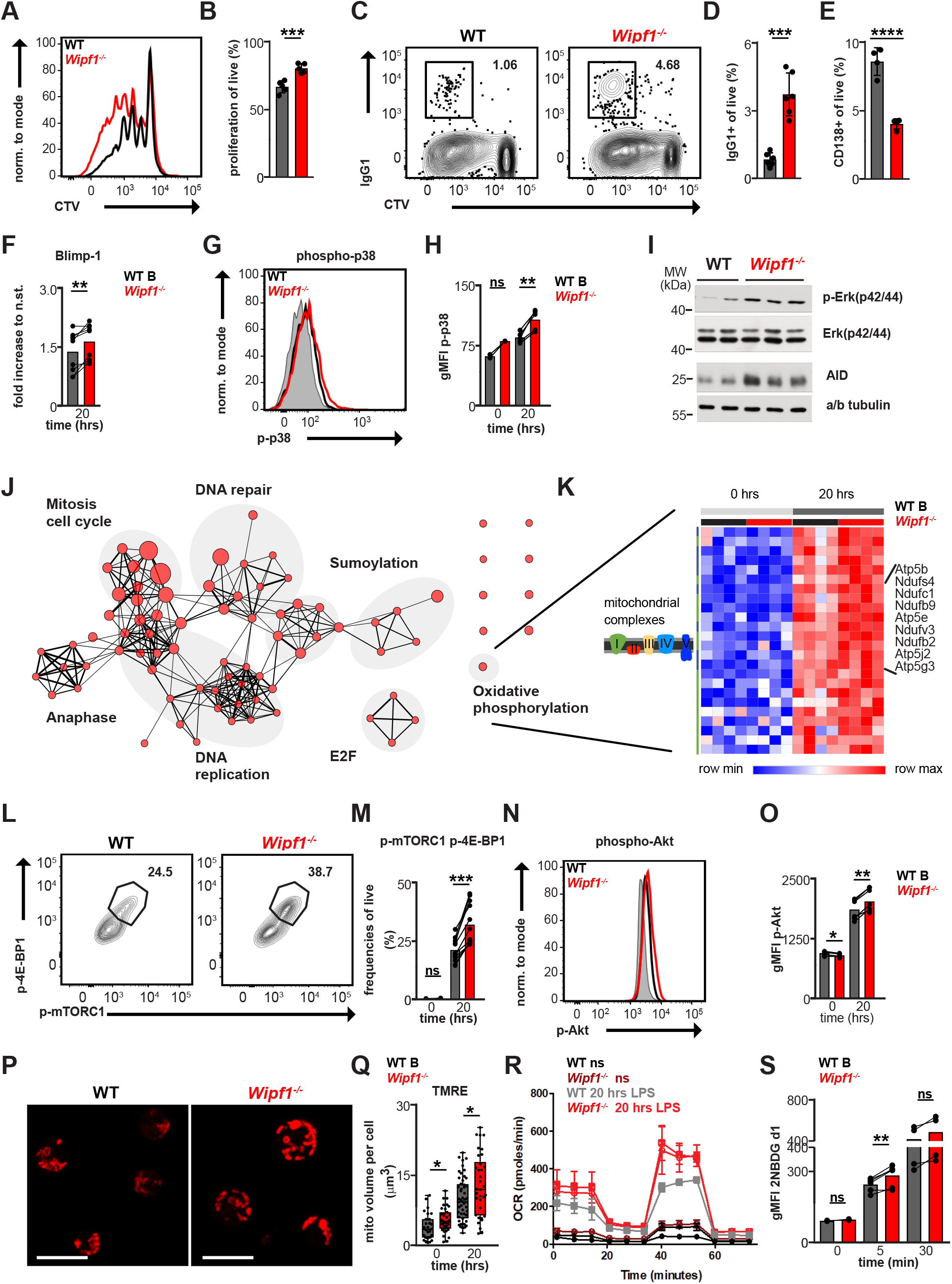
WIP deficiency metabolically primes B cells for PC differentiation. **(A-E)** CTV labeled splenic WT or *Wipf1*^*-/-*^ B cells were stimulated with LPS for 4 days. Flow cytometry measurements of **(A)** proliferation (CTV dilution), quantified in **(B)** and **(C)** class-switch recombination to IgG1, quantified in **(D)**, and CD138+ cells quantified in **(E)**. n≥4 mice per genotype (mean ± SEM). Statistical significance was calculated using student’s t test. **(F-S)** Splenic WT or *Wipf1*^*-/-*^ B cells were stimulated with LPS for 20 hrs. **(F-H)** Flow cytometry analysis of intracellular expression of **(F)** Blimp-1 **(G+H)** phospho-p38, quantified by analyzing the geometric mean fluorescent intensity (gMFI). n≥4 mice per genotype, pooled from multiple experiments. Statistical significance was calculated using paired student’s t test. **(I)** Immunoblot of stimulated B cells probed with antibodies as indicated. Data are representative of at least two independent experiments. **(J)** Gene Set Enrichment Analysis (GSEA). Nodes indicate significantly altered gene sets (FDR < 0.05). Node size indicates the size of each gene set. Red node color indicates up-regulation (normalized enrichment score > 0). Groups of similar gene sets were annotated manually. Network was generated using EnrichmentMap. **(K)** Expression of genes encoding the five complexes of the electron transport chain (extract, Wiki Pathways Oxidative Phosphorylation). Color indicates z-scored DESeq2-normalized gene counts. Rows are grouped by complex and clustered hierarchically (1-pearson correlation, average linkage). n=4 mice per genotype. **(L+M)** Flow cytometry analysis of phosphorylation of 4E-BP1 and mTORC1 **(M)** Frequencies of cells demonstrating phosphorylation of 4E-BP1 as well as mTORC1 **(N)** phosphorylation of Akt, **(O)** quantified by analyzing the geometric mean fluorescent intensity (gMFI). n≥4 mice per genotype, pooled from multiple experiments. Statistical significance was calculated using paired student’s t test. **(P)** B cells were stained for TMRE and imaged by confocal microscopy (scale bar 10 μm) **(Q)** Quantification of the volume of TMRE positive mitochondria per cell. Each dot indicates one cell. **(R)** Oxygen consumption rate (OCR) was measured in technical triplicates of B cells isolated from 1 WT mouse and 2 *Wipf1*^*-/-*^ mice using Seahorse flux technology. One of 2 experiments is shown (n=4 per genotype in total, mean ± SD). **(S)** Cells were incubated with 2NBDG for 5 and 30 min, uptake determined using flow cytometry and quantified by analyzing the geometric mean fluorescent intensity (gMFI). n≥4 mice per genotype, pooled from multiple experiments. Statistical significance was calculated using paired student’s t test. *p < 0.05, **p < 0.01, ***p < 0.001, ****p < 0.0001, ns = not significant

We next characterized CD4 T cell subsets in the colonic LP of *Wipf1*^*-/-*^ mice by re-clustering of the mass cytometric dataset shown in Fig. 1 (Fig. 2D+E). We found an increased population of activated CD4 T cells isolated from the colonic LP of *Wipf1*^*-/-*^ mice, with about 70% of CD4 T cells expressing CD44 compared to about 30% of CD4 T cells isolated from WT colonic LP (Fig. 2F). Of these activated CD4 T cells, 45% of *Wipf1*^*-/-*^ CD4 T cells expressed the transcription factor IRF4 compared to 20% of WT CD4 T cells, indicating a TfH cell phenotype (Fig. 2G). The frequency of FoxP3 expressing, regulatory CD4 T cells was similar between WT and *Wipf1*^*-/-*^ mice (Fig. 2G). Thus, in line with enhanced follicles and GC B cell responses, TfH cell numbers were increased in the colon of *Wipf1*^*-/-*^ mice as compared to controls.

Concomitant with the enhanced TD immune response, we observed a significant increase in the frequencies and numbers of IgA and IgG1 PC in the colon and mLN of *Wipf1*^*-/-*^ mice (Fig. 2H+I, Fig. S2G+H), as well as enhanced intestinal IgA, IgG1 and IgM antibodies (Fig. 2J, S2I). Notably, we detected significantly elevated levels of IgA and IgG1 but also other isotypes in the sera of non-immunized *Wipf1*^*-/-*^ mice but not in WT littermate controls (Fig. S2J). Strikingly, we found serum antibodies of *Wipf1*^*-/-*^ mice to be auto-reactive against single- and double stranded DNA as well as cardiolipin (Fig. 2K). Of note, serum levels of auto-antibodies were reduced in *Wipf1*^*-/-*^ mice treated with antibiotics to levels in sera of WT controls (Fig. 2K). This data hence suggest that *Wipf1*^*-/-*^ B cells in the inflamed colonic LP overreact to the microbiome, leading to the production of systemic, microbe-dependent, autoantibodies.

The intrinsic inflammatory activity of autoantibodies is dependent on the glycosylation pattern, which determines the properties of Fc-receptor binding and hence the activation of monocytes. In sera of human IBD patients, hypo-sialylated IgG has been described (*20*). To test whether aberrantly produced and glycosylated IgG contributes to inflammatory monocyte activation, we next measured the sialylation of IgG purified from sera of *Wipf1*^*-/-*^ mice by ELISA using biotinylated Sambucus nigra, a lectin specific for terminal-linked (2,6)-sialic acid (*21*). We found that, indeed, *Wipf1*^*-/-*^ IgG demonstrate lower levels of sialylation compared to WT IgG (Fig. 2L). We next stimulated WT bone-marrow derived macrophages (BMDMs) with purified IgG from WT or *Wipf1*^*-/-*^ mice. Hypo-sialylated *Wipf1*^*-/-*^ IgG enhanced the relative expression levels of *Nlrp3* and *Thioredoxin* mRNA in BMDMs (Fig. 2M). Thioredoxin has been described to promote NLRP3 inflammasome activation and IL-1β production in BMDMs (*22*). Indeed, we detected higher concentrations of IL-1β as well as IFN-γ in the supernatants of BMDMs treated with hypo-sialylated *Wipf1*^*-/-*^ IgG (Fig. 2N+O). We concluded that hypo-sialylated IgG produced during inflammation in an autoimmune setting in *Wipf1*^*-/-*^ mice supports inflammatory monocyte activation.

Together these findings suggest an ongoing TD immune response in the colonic LP of *Wipf1*^*-/-*^ mice concomitant with an aberrant production of intestinal IgG and a microbiome-dependent production of systemic auto-reactive antibodies. Our results further provide evidence that intestinal inflammation might enhance the intrinsic inflammatory activity of IgG which has the potential to exacerbate inflammation through monocyte/macrophage activation.

### WIP-deficiency metabolically primes B cells for PC differentiation

Concurrent with the enhanced microbiome-dependent serum auto-antibodies in *Wipf1*^*-/-*^ mice, stimulating purified B cells with LPS *in vitro*, we found a significantly increased frequency of proliferating *Wipf1*^*-/-*^ B cells (Fig. 3A+B) as well as increased CSR and decreased expression of the PC marker CD138 (*16*). One essential transcription factor for initiating and driving ASC generation is Blimp-1, which is induced by signaling pathways including Erk-kinases (*23*). 20 hours after LPS stimulation survival and blasting of B cells were similar between WT and *Wipf1*^*-/-*^ B cells (Fig. S3A+B). We found a stronger up-regulation of Blimp-1 in *Wipf1*^*-/-*^ B cells using flow cytometry when normalized on the expression levels of ex vivo isolated lymphocytes (Fig. 3F, Fig. S3C). Concomitantly, we found enhanced phosphorylation of Erk as well as p38 in *Wipf1*^*-/-*^ B cells, indicating an enhanced activation of MAPK pathways (Fig. 3G-I, Fig. S3D). CSR is dependent on the B cell–specific enzyme activation-induced cytidine deaminase (AID; (*24, 25*)). AID expression was enhanced in *Wipf1*^*-/-*^ B cells 20 hrs after LPS stimulation (Fig. 3I, Fig. S3E). Collectively, our data suggests that enhanced activation of MAPK pathways together with Blimp-1 and AID expression might provide the molecular basis for the efficient CSR and differentiation of *Wipf1*^*-/-*^ B cells into ASC *in vitro* and *in vivo*.

We next performed bulk RNA sequencing of WT and *Wipf1*^*-/-*^ B cells *ex vivo* or stimulated for 20 hours with LPS *in vitro*. Gene set enrichment analyses (GSEA) of *Wipf1*^*-/-*^ transcriptomes demonstrated enrichments of gene transcripts associated with Mitosis, DNA replication and DNA repair, but also oxidative phosphorylation (OxPhos) (KEGG and HALLMARK genesets) (Fig. 3J). Especially transcripts of genes associated with the mitochondrial complexes I – V were enriched in *Wipf1*^*-/-*^ B cells (Fig. 3K, Fig. S3F). The mammalian target of rapamycin (mTORC1) complex is known to drive mitochondrial biogenesis by promoting translation of nucleus-encoded mitochondria-related mRNAs of complex I-V via inhibition of the eukaryotic translation initiation factor 4E binding proteins (4E-BP1) (*26*). Hyperphosphorylation of 4E-BP1 by mTORC1 as well as the PI3K/Akt pathway results in activation of cap-dependent translation (*27-29*). We next assessed the phosphorylation of mTOR, pAkt and 4E-BP1 by flow cytometry. We found that mTOR as well as 4E-BP1 were robustly phosphorylated in WT and *Wipf1*^*-/-*^ B cells (Fig. S3G+H). Intriguingly, we noticed a 2-fold increased frequency of *Wipf1*^*-/-*^ B cells showing enhanced phosphorylation of both, mTORC1 and 4E-BP1 (Fig. 3L+M). Similarly, *Wipf1*^*-/-*^ B cells demonstrated an enhanced Akt phosphorylation compared to WT B cells, albeit lower basal Akt phosphorylation when analyzed ex vivo (Fig. 3N+O, Fig. S3I+J). Together, this data suggests that efficient activation of the mTOR/Akt/4E-BP1 pathway 20 hours after LPS stimulation might heighten mitochondrial biogenesis in *Wipf1*^*-/-*^ B cells. Indeed, using tetramethylrhodamine ethyl ester (TMRE) staining, we found the volume of mitochondria in *Wipf1*^*-/-*^ B cells increased compared to control B cells (Fig. 3P+Q, Fig. S3K). We next analyzed mitochondrial respiration using the Seahorse assay. *Wipf1*^*-/-*^ B cells demonstrated a slightly increased basal respiratory capacity and a significantly enhanced maximal oxygen consumption rate compared to WT B cells 20 hrs after LPS stimulation (Fig. 3R). Additionally, LPS-stimulated *Wipf1*^*-/-*^ B cells demonstrated enhanced uptake of the fluorescent glucose analogue 2-(N-(7-nitrobenz-2-oxa-1,3-diazol-4-yl)amino)-2-deoxyglucose (2-NBDG) after 5 min of incubation (Fig. 3S).

Together, the absence of WIP in B cells enhanced the MAPK/Erk and mTOR/Akt/4E-BP1 pathway 20 hours after LPS stimulation leading to a state of heightened metabolic activity.

### *Wipf1*^*-/-*^ B cells boost pro-inflammatory cytokine secretion by intestinal CD4 T cells

Pro-inflammatory cytokines - such as IL-1β, IL-6, Tumour necrosis factor alpha (TNF-α), IL-17 and Interferon-γ (IFN-γ), have been directly implicated in the pathogenesis of IBD, controlling intestinal inflammation and the associated clinical symptoms (*4*). We next assessed the production of these cytokines by intracellular staining and flow cytometry of re-stimulated colonic CD4 T cells. We found an enhanced frequency of cytokine-producing CD4 T cells isolated from *Wipf1*^*-/-*^ mice (Fig. 4 A+B, Fig. S4A). Similarly, we detected more CD44^+^ activated CD4 T cells (Fig. S4B) producing pro-inflammatory cytokines (Fig. S4C) in the intestinal draining mLN of *Wipf1*^*-/-*^ mice compared to their WT counterparts. Of note, CD4 T cells co-expressing IFN-γ and GM-CSF or IL-17A and IL-17F were detectable only in the colonic LP or mLN of *Wipf1*^*-/-*^ mice (Fig. 4 A+B, Fig. S4A). Together these findings showed that CD4 T cells produce colitis-associated inflammatory cytokines characteristic for the pro-inflammatory TH1/TH17 T cell subsets in the inflamed colonic tissue of *Wipf1*^*-/-*^ mice.

**Fig. 4:**
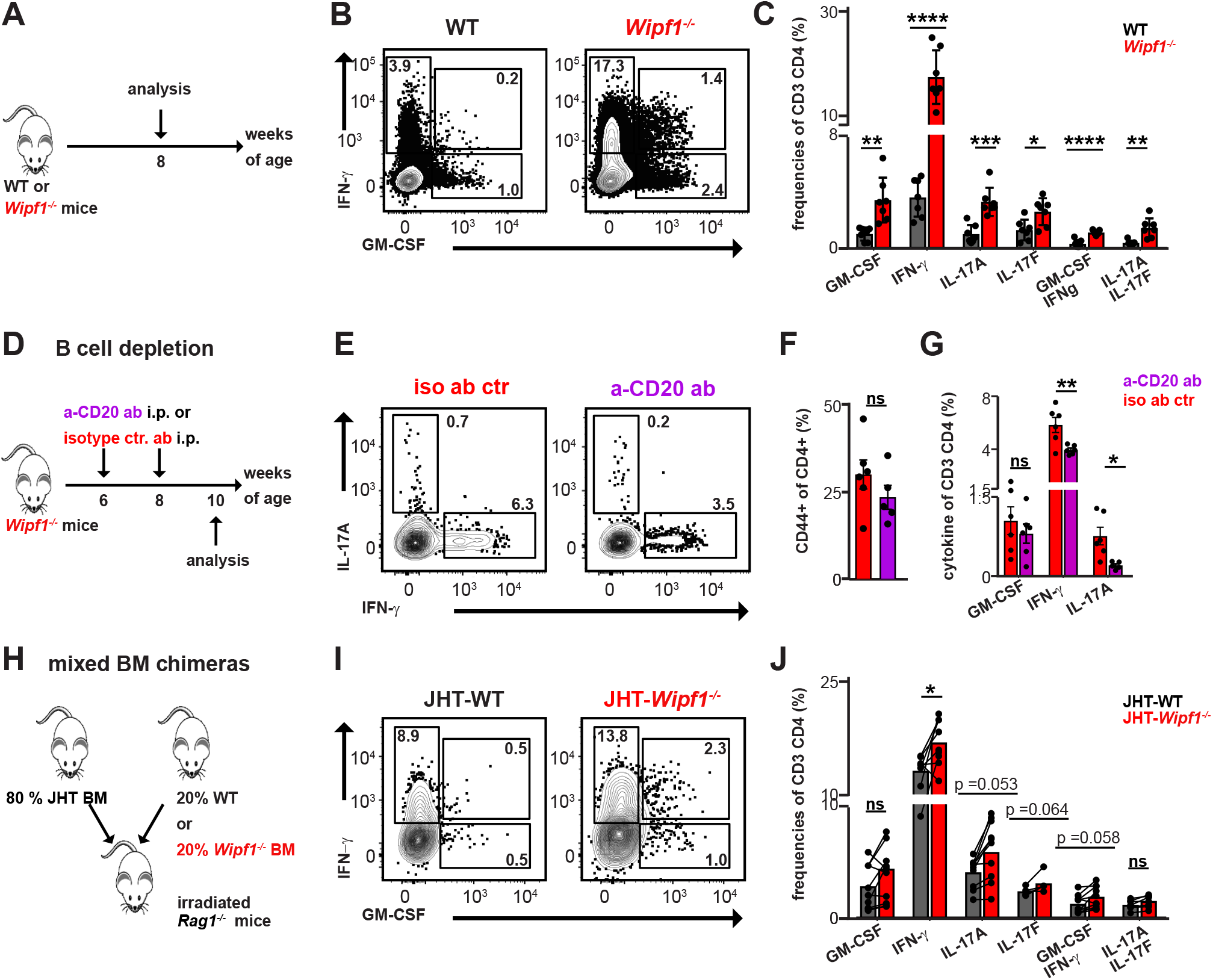
*Wipf1*^*-/-*^ B cells boost pro-inflammatory cytokine secretion by intestinal CD4 T cells. **(A)** Schematics indicating the time-point of analysis of cytokine production of CD4 T cells isolated from colonic tissue of WT or *Wipf1*^*-/-*^ mice. **(B+C, E-G, I+J)** Lymphocytes were isolated from **(B+C, I+J)** colonic tissue or **(E-G)** mLN of **(B+C)** WT or *Wipf1*^*-/-*^ mice, **(E-G)** *Wipf1*^*-/-*^ mice treated with either isotype control antibody or anti-CD20 antibody to deplete B cells, or **(I+J)** JHT-WT or JHT-*Wipf1*^*-/-*^ mixed BM chimeric mice and stimulated with Brefeldin A and PMA/Ionomycin for 4 hours. **(B, E, I)** Representative flow cytometric dot blots of CD4 T cells intracellularly expressing indicated cytokines. **(C, F, J)** Frequencies of intracellular staining of indicated cytokines of CD4 T cells after in vitro re-stimulation. **(D)** Experimental setup of B cell depletion in *Wipf1*^*-/-*^ mice. **(G)** Frequencies of CD44^+^ CD4 T cells. **(H)** Experimental setup of the generation of JHT-WT or JHT-*Wipf1*^*-/-*^ mixed bone-marrow (BM) chimeric mice. **(A-J)** n≥6 mice per genotype, pooled from 2-3 independent experiments. Statistical significance was calculated using student’s t test. *p < 0.05, **p < 0.01, ***p < 0.001, ****p < 0.0001, ns = not significant

To assess the role of B cells in driving CD4 T cell cytokine production, we depleted B cells of *Wipf1*^*-/-*^ mice at 6 and 8 weeks of age and hence before the manifestation of colonic inflammation using an anti-CD20 antibody (Fig. 4D). B cells were efficiently depleted in the spleen and mLN (Fig. S4D), which resulted in a 50% reduction of total lymphocyte numbers in mLN of *Wipf1*^*-/-*^ mice (Fig. S4E). We found incomplete depletion of B cells in PP as well as the peritoneal cavity (Fig. S4D). Anti-CD20 treatment of *Wipf1*^*-/-*^ mice led to a reduction in IgA as well as IgG1 PC (Fig. S4F). Concurrent with the reduced ASCs, serum autoantibodies (Fig. S4G) as well as intestinal antibody levels in colonic mucus (Fig. S4H) were reduced in anti-CD20 treated *Wipf1*^*-/-*^ mice. Notably, B cell depletion in *Wipf1*^*-/-*^ mice did not influence the frequencies of CD4 T cells isolated from mLN (Fig. S4I), but led to diminished frequencies of CD44^+^ CD4 T cells as well as reduced intracellular IFN-γ and IL-17A staining (Fig. 4D-G) in CD4 T cells isolated from mLN. Thus, B cell depletion during the onset of intestinal inflammation reduced CD4 T cell activation and cytokine production.

To establish whether the absence of WIP exclusively in B cells has an effect on pro-inflammatory cytokine production of colonic T cells, we next generated mixed BM chimeras by adoptively transferring a mix of BM from either WT or *Wipf1*^*-/-*^ mice and BM from mice that lack B cells (JHT mice) into lethally irradiated *Rag1*^*-/-*^ recipients (Fig. 4H). In this setting, all newly generated B cell will be *Wipf1*^*-/-*^ in an environment containing mainly WT cells. 10 weeks after adoptive transfer, B cells have repopulated the colonic tissue with a slight reduction in B cell (Fig. S4J) as well as IgA PC frequencies (Fig. S4K) in JHT-*Wipf1*^*-/-*^ chimeric mice compared to controls. Nevertheless, we found increased IgA antibody levels in the mucus and enhanced levels of serum IgM in JHT-*Wipf1*^*-/-*^ chimeric mice compared to controls (Fig. S4L+M). Strikingly, we detected a pronounced increased frequency of activated CD44^+^ CD4 T cells (Fig. S4N) concomitant with enhanced pro-inflammatory cytokine production of CD4 T cells (GM-CSF, IFN-γ, IL-17A and IL-17F) isolated from the LP of the inflamed colon of JHT-*Wipf1*^*-/-*^ chimeric mice (Fig. 4I+J), albeit less pronounced as in complete *Wipf1*^*-/-*^ mice.

Collectively, our data provides evidence that *Wipf1*^*-/-*^ B cells directly promote pro-inflammatory cytokine production of CD4 T cells, thus likely enhancing intestinal inflammation.

### *Wipf1*^*-/-*^ B cells are sufficient to enhance pro-inflammatory cytokine production in WT CD4 T cells in an acute model of colitis

To address the influence of B cells on IBD development in an acute model of colitis, in which again, only B cells lack WIP, we made use of a T cell transfer colitis model. In the absence of Tregs, the transfer of naïve CD4^+^CD45RB^high^ WT T cells generates large numbers of disease-promoting TH1 and TH17 cells (*30*). We co-transferred purified B cells isolated from mLN of WT or *Wipf1*^*-/-*^ mice together with naïve WT T cells into *Rag1*^*-/-*^ recipients and analysed lymphocyte populations isolated from colonic tissue or mLN 8 weeks after transfer (Fig. 5A). Weekly scoring of the mice showed a weight gain of *Rag1*^*-/-*^ recipients which received WT B cells as compared to no B cell transfer, which was reduced if *Wipf1*^*-/-*^ B cells were transferred (Fig. 5B). Interestingly, transfer of *Wipf1*^*-/-*^ B cells led to an enhanced colonic weight to length ratio concomitant with increased total cell numbers isolated from mLN or colonic tissue of *Rag1*^*-/-*^ recipients, compared to controls (Fig. 5C+D). We noticed that most of the transferred *Wipf1*^*-/-*^ B cells differentiated into IgA ASCs, with only a modest reduction in total cell numbers and frequencies of IRF4^+^ IgA^+^ CD19^low^ cells isolated from colonic tissue (Fig. 5E-G, Fig. S5A-C). This decrease was reflected in reduced levels of IgA antibodies detected in the mucus of recipients which received *Wipf1*^*-/-*^ B cell (Fig. S5D). Concurrent with the increase in total cell numbers in mLN in *Wipf1*^*-/-*^ B cell co-transferred recipients, numbers and frequencies of total CD4 T cells as well as activated CD44^+^ CD4 T cells were increased in those mLN but not in lymphocyte populations isolated from the colon (Fig. S5E-H). Strikingly, CD4 T cells isolated from mLN of *Wipf1*^*-/-*^ B cell transferred mice demonstrated significantly enhanced numbers and a modest increase in frequencies of pro-inflammatory cytokine expressing cells (Fig. 5H+I, Fig. S5K-O). This was less apparent in CD4 T cells isolated from colonic tissue, most likely due to the strong acute inflammation in the colons of all mice. These results again reinforce a role for *Wipf1*^*-/-*^ B cells in fueling intestinal inflammation through CD4 T cell activation.

**Figure 5:**
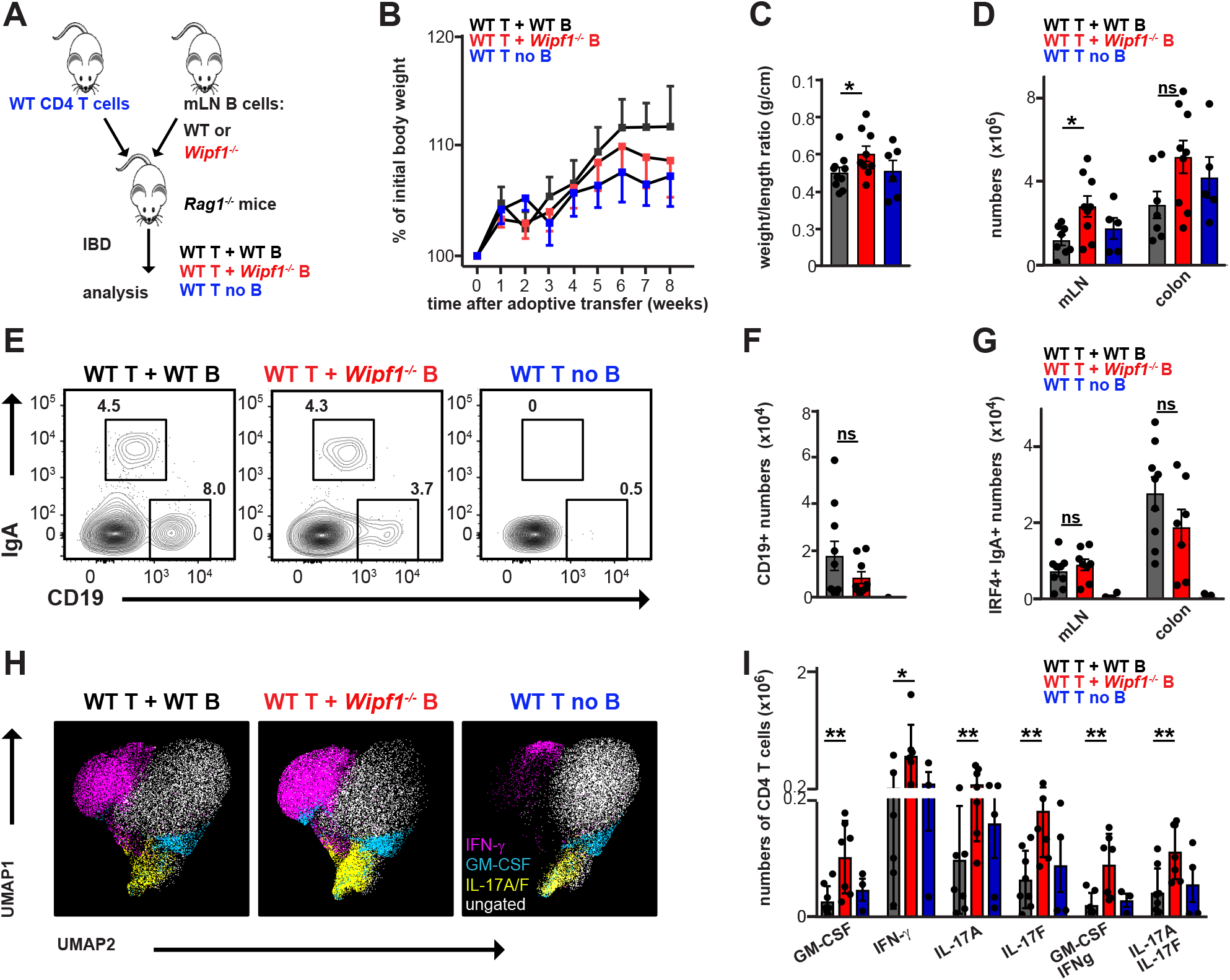
*Wipf1*^*-/-*^ B cells are sufficient to enhance pro-inflammatory cytokine production in WT CD4 T cells in an acute model of colitis. **(A)** Schematics indicating the experimental setup of the adoptive transfer of naïve T cells together with WT or *Wipf1*^*-/-*^ B cells into *Rag1*^*-/-*^ recipients to induce colitis. **(B)** Body weights shown as percentage of starting weight. **(C)** Length of the proximal colon as well as the weight was measured and weight to length ratio calculated. Statistical significance was calculated using Mann-Whitney U-test. **(D)** Total cell numbers isolated from mLNs or colonic tissue. **(E)** Representative flow cytometric dot blots of B cells (CD19^+^) and IgA PC (IgA^+^,CD19^low^) isolated from mLNs. **(F)** Total numbers of B cells isolated from mLNs. **(G)** Total numbers of IgA^+^ PC isolated from mLNs or colonic tissue. **(H+I)** Lymphocytes were isolated from mLNs and stimulated with Brefeldin A and PMA/Ionomycin for 4 hours. **(H)** UMAP was used to depict cytokine producing CD4 T cells. Manually gated CD4 T cells expressing indicated cytokines are overlaid as color dimension. **(I)** Total cell numbers of CD4 T cells isolated from mLN expressing indicated cytokines intracellularly after in vitro re-stimulation. **(A-I)** n≥5 per genotype, pooled from 3 independent experiments (all mean+/-SEM). Statistical significance was calculated using student’s t test unless otherwise indicated. *p < 0.05, **p < 0.01, ***p < 0.001, ****p < 0.0001, ns = not significant

### B cell mediated co-stimulation via CD86 enhances CD4 T cell cytokine production

To determine the effect of B cell on T cell activation in a more controlled environment, we used a mixed-lymphocyte reaction to activate T cells by means of MHC-II mismatch. We stimulated purified mLN B cells from WT or *Wipf1*^*-/-*^ Balb/c mice with LPS and cultured them with naïve, splenic CD4 T isolated from C57BL/6 mice (Fig.6A). We found GM-CSF and IFN-γ the most prevalent cytokines secreted into the supernatants of those co-cultures Fig. 6B). B cells contributed to IL-6 and IL-10 cytokine levels in the culture supernatant, and *Wipf1*^*-/-*^ B cells were more efficient in secreting those cytokines compared to WT B cells (Fig. S6A). The concentration of both, GM-CSF and IFN-γ, but also TNF-a, IL-6 and IL-10 were significantly increased in the co-culture supernatants of WT T/*Wipf1*^*-/-*^ B cells compared to supernatants of WT T/WT B cell co-cultures (Fig. 6B). Thus, the presence of *Wipf1*^*-/-*^ B cells enhanced the secretion of GM-CSF as well as IFN-γ of WT CD4 T cells *in vitro* as well as *in vivo*.

**Figure 6:**
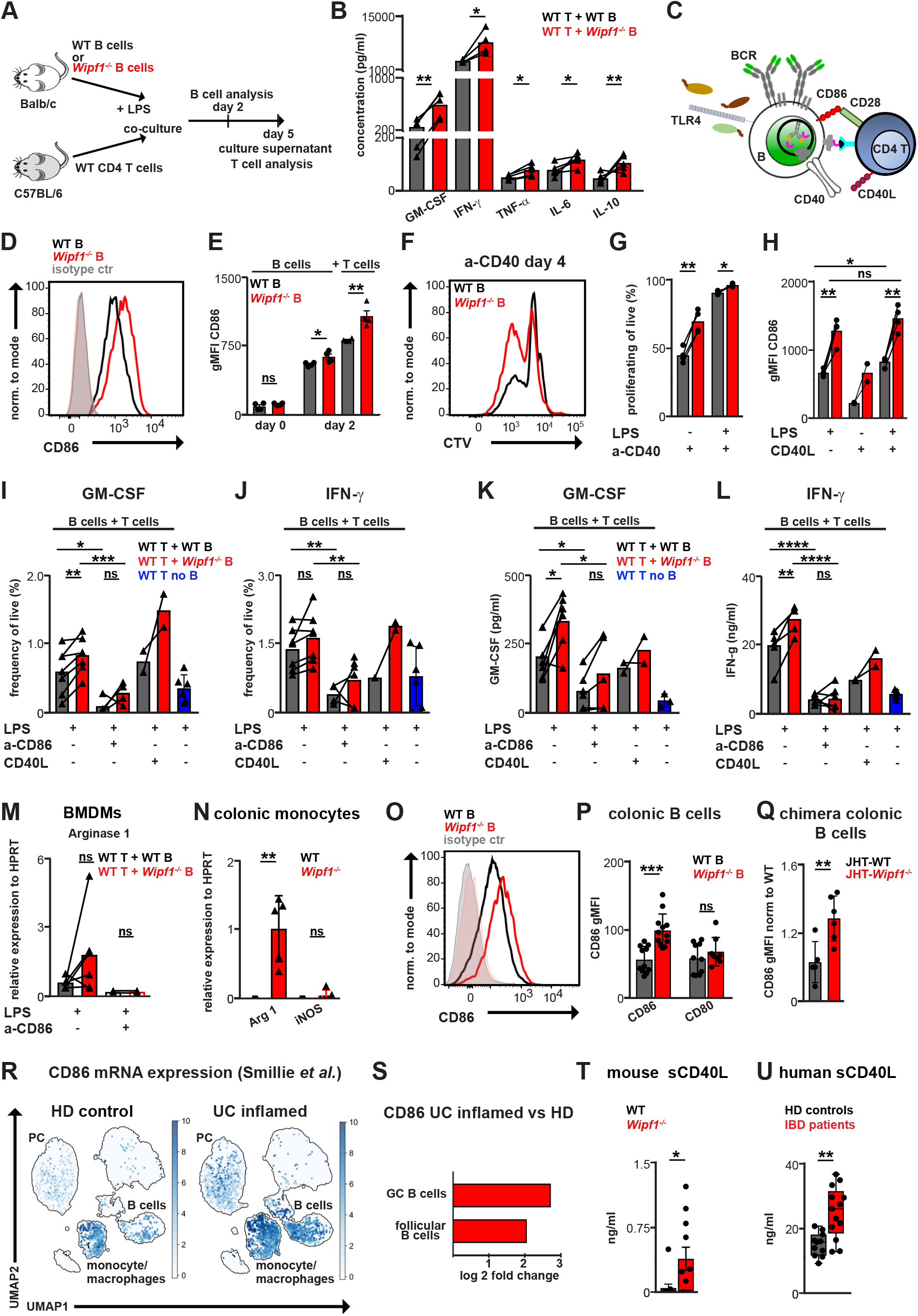
B cell mediated co-stimulation via CD86 enhances CD4 T cell cytokine production. **(A)** Schematics indicating the experimental setup of the mixed-lymphocyte reaction. **(B)** Indicated cytokines secreted into the co-culture supernatants were analyzed using a multiplex assay. **(C)** Schematics indicating surface molecules involved in B/T interactions analyzed. **(D+E)** Flow cytometry analysis of CD86 expression on mLN WT or *Wipf1*^*-/-*^ B cells stimulated with LPS for 48 hrs in the presence of T cells **(E)** CD86 expression quantified by analyzing the geometric mean fluorescent intensity (gMFI). **(B, D+E)** n≥4 mice per genotype, pooled from multiple experiments. **(F+G)** CTV labeled mLN WT or *Wipf1*^*-/-*^ B cells were stimulated with anti-CD40 for 4 days. Flow cytometry measurements of **(F)** proliferation (CTV dilution), quantified in **(G). (H)** Flow cytometry analysis of CD86 expression on B cells stimulated for 48 hrs with LPS, CD40L or both and quantified by analyzing the gMFI. **(F-H)** n≥3 mice per genotype (except CD40L alone n=2), pooled from multiple experiments. **(I-J)** Frequencies of **(I)** GM-CSF or **(J)** IFN-γ expressing CD4 T cells stimulated with Brefeldin A and PMA/Ionomycin for 4 hours, stained and analyzed by flow cytometry at day 5 of co-culture. **(K+L)** Indicated cytokines secreted into the co-culture supernatants in the presence or absence of indicated stimuli were analyzed using a multiplex assay. n≥4 mice per genotype (except CD40L+LPS stimulation n=2), pooled from multiple experiments. **(M)** BMDMs were incubated with indicated co-culture supernatants (n=5 experiments) for 20 hrs. Relative mRNA expression of *Arginase-1* normalized to HPRT determined by qPCR **(N)** Relative mRNA expression of *Arginase-1* and *iNOS* normalized to HPRT of sorted CD11b^+^ ex vivo isolated colonic monocytes determined by qPCR. n=5 mice per genotype. **(O-Q)** CD86 expression shown as gMFI on ex vivo isolated colonic B cells of **(O+P)** WT or *Wipf1*^*-/-*^ mice or **(Q)** JHT-WT or JHT-*Wipf1*^*-/-*^ mixed BM chimeric mice analyzed by flow cytometry. n≥5 mice per genotype. **(R+S)** Single-cell RNA sequencing (scRNA-seq) analysis demonstrates significant upregulation of CD86 in colonic biopsies of inflamed UC patients compared to healthy tissue (data extracted from *33*). **(T+U)** sCD40L detected in sera of **(T)** WT or *Wipf1*^*-/-*^ mice (n≥10 mice per genotype) or **(U)** healthy donors (HD) or IBD patients (n≥10 sera per group). Statistical significance was calculated using **(B,G+H, I-M)** paired student’s t test or **(E, N-P, R+S)** student’s t test. *p < 0.05, **p < 0.01, ***p < 0.001, ****p < 0.0001, ns = not significant

In addition to the mismatched MHC-II/TCR activation, B cells interact with T cells by means of co-stimulatory molecules such as CD86/CD80 engaging CD28 or CD40 engaging CD40L (Fig. 6C). We found that, compared to WT B cells, *Wipf1*^*-/-*^ B cells expressed higher levels of CD86 on their surface, which is further increased by the presence of T cells (Fig. 6D+E, Fig. S6B+C). In contrast, expression levels of CD80, CD40 and MHC-II at day 2 after LPS stimulation in the presence or absence of T cells were similar between WT and *Wipf1*^*-/-*^ B cells (Fig. S6D-I). Notably *Wipf1*^*-/-*^ B cells isolated from JHT-*Wipf1*^*-/-*^ chimeric mice and thus from a non-inflamed environment demonstrated enhanced CD86 expression compared to JHT-WT B cells in the presence of T cells, indicating an intrinsic effect of *Wipf1*^*-/-*^ B cells (Fig. S6J). We noted a slightly higher expression of CD40 on ex vivo isolated *Wipf1*^*-/-*^ B cells (Fig. S6G). Stimulating B cells with an activating a-CD40 antibody led to an enhanced proliferation of *Wipf1*^*-/-*^ B cells compared to WT B cells (Fig. 6F+G), which was further increased by adding additional LPS (Fig. 6G). Of note, the expression level of CD86 was boosted further on WT and *Wipf1*^*-/-*^ B cells by the combined stimulation with CD40L and LPS (Fig. 6H).

Next, we sought to determine the role of CD86 for CD4 T cell cytokine production using anti-CD86 blocking antibodies. Co-culturing WT CD4 T with *Wipf1*^*-/-*^ B cells elevated the frequencies of cytokine producing CD4 T cells as well as secretion of GM-CSF and IFN-γ compared to the presence of WT B cells (Fig. 6I-L, Fig. S6K+L). This was not due to an increase in cell numbers of *Wipf1*^*-/-*^ B cells, as, despite enhanced proliferation of *Wipf1*^*-/-*^ B cells after LPS stimulation, frequencies were comparable to WT B cells at days 2 and 5 of co-culture (Fig. S6M), likely due to the previously reported diminished survival of *Wipf1*^*-/-*^ B cells (*16*). Notably, co-culturing WT CD4 T with JHT-*Wipf1*^*-/-*^ B cells elevated the frequencies of GM-CSF producing CD4 T cells compared to the presence of JHT-WT B cells (Fig. S6N), again indicating an intrinsic effect of *Wipf1*^*-/-*^ B cells. Blocking B/T interaction by anti-CD86 antibodies reduced the frequencies of cytokine producing CD4 T cells as well as secretion of GM-CSF and IFN-γ (Fig. 6I-L). Adding additional CD40L into the culture medium further enhanced frequencies of GM-CSF but not IFN-γ expressing CD4 T cells (Fig. 6I+J), but did not impact on GM-CSF or IFN-γ secretion into the supernatant (Fig.6 K+L). Collectively, these findings suggest that cell intrinsic hyper-reactivity of *Wipf1*^*-/-*^ B cells towards LPS and CD40L stimulation enhances CD86 expression and hence the co-stimulatory capacity of *Wipf1*^*-/-*^ B cells, thereby exacerbating CD4 T cell pro-inflammatory cytokine production.

T cell derived cytokines, such as GM-CSF, might enhance the recruitment and the differentiation of inflammatory monocytes (*31*). BMDMs cultured with supernatant of WT T/*Wipf1*^*-/-*^ B cells demonstrated higher relative mRNA expression levels of Arginase-1 (Fig. 6M), but not iNOS (Fig. S6P) compared to supernatant of WT T/WT B cells. Notably, expression of mRNA was abrogated when co-cultures were treated with a-CD86 antibody (Fig. 6M) and hence contained reduced levels of GM-CSF and IFN-γ. Of note, *ex vivo* isolated, colonic CD11b^+^ monocyte/macrophages from *Wipf1*^*-/-*^ mice similarly expressed enhanced relative mRNA expression levels of Arginase-1 but not iNOS compared to WT controls, indicating a pro-fibrotic phenotype. Thus, B cell-induced T cell cytokines might help to differentiate inflammatory, pro-fibrotic monocytes.

Consistent with our *in vitro* findings, we observed a higher expression of CD86, but not CD80 on *Wipf1*^*-/-*^ B cells compared to WT B cells isolated from the colonic LP (Fig. 6O+P). Again, B cells isolated from colonic tissue of JHT-*Wipf1*^*-/-*^ chimeric mice and thus in a non-inflamed environment also demonstrated enhanced CD86 expression (Fig. 6Q). The expression of both, CD86 and CD80, was similar or even reduced on monocytes isolated from colonic tissue of WT or *Wipf1*^*-/-*^ mice (Fig.S6Q), indicating a B cell specific effect. Of note, CD86 mRNA transcripts are significantly increased in bulk RNA sequencing data comparing colonic tissue of UC patients to healthy individuals (Fig. S6R) (*32*). Interestingly, CD86 is amongst the top 100 significantly upregulated genes in GC and follicular B cells in a single-cell RNA sequencing (scRNA-seq) analysis of differentially expressed genes in colonic biopsies of inflamed UC patients compared to healthy tissue (Fig. 6R+S, Fig. S6S and (*33*)). Hence CD86 expression on B cells might push inflammatory cytokine production in CD4 T cells also in IBD patients. Our *in vitro* data implied a role for CD40 stimulation by CD40L in boosting CD86 expression on B cells. We indeed found sCD40L to be elevated in both, sera of *Wipf1*^*-/-*^ mice as well as IBD patients compared to controls (Fig.6 T+U).

Collectively the data presented here suggest that up-regulation of the co-stimulatory molecule CD86 might play a crucial role in priming pro-inflammatory cytokine production in CD4 T cells in the inflamed colonic tissue. Enhanced expression of CD86 on B cells could be boosted by the presence of sCD40L in our mouse model of chronic colitis but also in human IBD patients.

## DISCUSSION

Perturbations in humoral immunity have been described during IBD, including the accumulation of intestinal B cells, an IBD specific subset of TfH cells and IgG PC in mucosal tissue (*8, 9*). Our study sheds light on the mechanism underpinning these observations, demonstrating a crucial role for B cells during autoimmune intestinal inflammation. Our findings suggest that during intestinal inflammation, microbial components and likely the general inflammatory milieu trigger enhanced expression of co-stimulatory CD86 on B cells. Increased CD86 then mediates pro-inflammatory cytokine production of T cells, which themselves provide co-stimulation to the B cells for improved activation and differentiation into ASC (IgA and IgG). B cell activation and co-stimulatory molecule expression can be helped additionally by sCD40L present during IBD. Together, this will eventually lead to the production of inflammatory IgG. B cell induced T cell cytokines, especially IFN-γ and GM-CSF might help to attract Ly6C^+^ inflammatory monocytes, while hypo-sialylated, inflammatory IgG secreted by differentiated ASC further activates those inflammatory monocytes to increase intestinal inflammation. We identify B cells as modulators of autoimmune IBD pathogenesis and suggest that our mouse model could be of immense value to test treatment strategies targeting chronic IBD.

Investigating *Wipf1*^*-/-*^ mice, we found that changes in the fecal microbiome correlated with the onset of disease. While younger *Wipf1*^*-/-*^ mice demonstrating bacterial diversity similar to that of WT mice, only two weeks older *Wipf1*^*-/-*^ mice demonstrated a reduced microbial diversity characteristic for intestinal inflammation. We found relatively reduced levels of *Lachnospiraceae*, similar to what has been described in CD patients (*34*). *Lachnospiraceae* are known to provide short-chain fatty acids (SCFA), microbial metabolites, which enhance the differentiation of regulatory B and T cells (*35, 36*). Interestingly, systemic autoreactive antibodies in the sera of *Wipf1*^*-/-*^ mice were dependent on the microbiome, as antibiotics treatment abrogated these antibodies. B cell receptor sequencing might identify if the altered composition of the microbiota or cross-reactivity to self-antigens plays a role in the production of systemic auto-reactive antibodies during IBD.

Our previous work demonstrated that WIP functions as a regulator of CD19 activation and PI3K/Akt signaling in B cells (*16*). WIP deficiency resulted in defects in B cell homing, chemotaxis, survival and differentiation, ultimately leading to diminished GC formation and antibody production after vaccination (*16*). At the same time, *Wipf1*^*-/-*^ mice demonstrate features of autoimmunity, such as autoantibody production (*14*), glomerulonephritis (*37*) and IBD similar to WAS patients. Mechanistically, we here described that hyper-reactivity of *Wipf1*^*-/-*^ B cells towards LPS stimulation led to a state of heightened metabolic activity mediated by efficient activation of the MAPK/Erk and mTOR/Akt/4E-BP1 pathways. These findings provide a likely cause for aberrant IgG production in colonic tissue, as well as for the systemic, microbe-dependent auto-antibodies in sera of *Wipf1*^*-/-*^ mice.

Although reduced after challenge vaccination (*16*), GC formation was enhanced in colonic tissue of *Wipf1*^*-/-*^ mice, indicating an ongoing TD immune response. B cell-dependent spontaneous GC formation concomitant with autoantibody production in spleen and lymph nodes during WAS has been linked to TLR7 and MyD88 signaling (*38, 39*) as well as B cell intrinsic IFN-γ signaling or IL-6 secretion (*40, 41*). Furthermore, B-cell intrinsic simultaneous deletion of CD80/CD86 abrogated GCs in WASp KO mice, leading to reduced serum autoantibodies (*42*). We here extent these findings by demonstrating that *Wipf1*^*-/-*^ B cells efficiently and specifically up-regulate CD86 after LPS stimulation, which enhanced co-stimulation of CD4 T cells and resulted in the exacerbated production of IFN-γ and GM-CSF. Of note, we detected enhanced CD86 expression on B cells, but not on monocytes, isolated from colonic tissue of *Wipf1*^*-/-*^ mice. CD86 up-regulation was independent of intestinal inflammation and hence an intrinsic feature of hyper-activated *Wipf1*^*-/-*^ B cells. Interestingly, recently published scRNAseq data comparing lymphocytes isolated from inflamed tissue biopsies of UC patients to tissue biopsies of HD found a significant increase of CD86 mRNA in GC and follicular B cells (*33*). Our data suggests that either genetic alterations (such as WIP deficiency) or likely the general inflammatory milieu in IBD patients triggers enhanced expression of the co-stimulatory molecule CD86 on B cells. Increased co-stimulation might then promote pro-inflammatory cytokine production of T cells. It was hypothesized that a UC specific IFN-γ expressing TfH cell subset could promote IgG CSR in active UC (*9*). Our data provides a possible explanation for these observations, demonstrating that B cells directly induce IFN-γ production by intestinal CD4 T cells.

In addition, we found that CD86 expression was enhanced by the presence of CD40. CD40L is directly supplied by CD4 T cells during B/T interaction and promotes CSR to IgG. Enhanced levels of sCD40L in the serum of *Wipf1*^*-/-*^ mice correlated with enhanced, microbial dependent serum (auto-)antibodies as well as the appearance of IgG1 plasmablast in the inflamed colonic tissue and IgG antibodies in the colonic mucus. Platelets have been described to express high levels of sCD40L in patients with WAS (*43*) and IBD (*44*), which correlated with the presence of autoantibodies as well as aberrant intestinal IgG production. Our data provides evidence that sCD40L locally elevates CD86 co-stimulation of B cells, thereby boosting CD4 T cell cytokine production and might additionally contribute to the aberrant production of intestinal IgG.

It has been speculated that similar to rheumatoid arthritis (RA) (*45*), changes in IgG glycosylation may contribute to enhanced monocyte activation during IBD. Indeed, hypo-sialylated IgG has been described in sera of human IBD patients (*20*). We found hypo-sialylated IgG present in the sera of our mouse model of chronic intestinal inflammation and demonstrated the capacity of this IgG to facilitate the differentiation of IL-1β secreting inflammatory BMDMs *in vitro*. It will be the interest of future studies to analyze the IgG glycome derived from intestinal ASC and hence present at the site of intestinal inflammation in mouse and human.

IgG mediated macrophage activation has been associated with intestinal inflammation in UC patients (*9, 10*). We found an enhanced accumulation of inflammatory monocytes in colonic tissue of *Wipf1*^*-/-*^ mice. Evidence from WASp^*-/-*^ mice suggests that inflammatory monocytes in the colonic LP are sufficient to promote intestinal inflammation (*46, 47*). Our data suggests an additional pathological mechanism, in which B cell-induced T cell cytokines (IFN-γ and GM-CSF) attract and promote differentiation of inflammatory, pro-fibrotic monocytes. In this line, recent findings described ILC-derived GM-CSF to promote the polarization of inflammatory macrophages during DSS induced colitis (*31*). During neuroinflammation, GM-CSF endorsed IL-1β production and phagocytosis, while IFN-γ initiated the differentiation of inflammatory monocytes (*48*). In addition to promoting GM-CSF and IFN-γ production, our results indicate that B cell-derived hypo-sialylated, inflammatory IgG further activates those inflammatory monocytes to escalate intestinal inflammation.

IBD therapy includes monoclonal antibody treatment against TNF-α and steroids; however, not all patients are responsive to these therapies. Failure to respond to anti-TNF-α treatment correlated with reduced *Lachnospiracea* taxa (*34*) but also with the presence of IgG PC and IgG-Fc complex activation signature of macrophages (*9, 10*). We here elucidate some of the mechanisms underpinning these observations and identify B cells, CD86 expression, sCD40L, IgG production and glycosylation as potential therapeutic targets.

Our data suggests that depleting B cells before the manifestation of disease and hence before the establishment of potentially inflammatory, IgG secreting ASC in intestinal tissues, has the potential to put a break on the vicious circle of intestinal inflammation. A small study analyzing B cell depletion in UC using rituximab (an anti-CD20 monoclonal antibody) has shown no benefit (*49*); but rituximab will not target potentially inflammatory PC in the intestinal tissue (*50*) and hence will not abrogate inflammatory IgG production. CD80/CD86 expression on B cells can be decreased by abatacept, the cytotoxic T lymphocyte antigen-4 (CTLA-4) immunoglobulin. Abatacept has been shown to diminish plamablasts and serum IgG in RA patients (*51*), but was not effective as treatment of CD or UC (*52*). It remains to be determined if blocking CD86 co-stimulation might be efficient in reducing inflammatory IgG antibodies and PC during IBD and the effect on intestinal inflammation. Of note, while anti-α4β7 integrin treatment might be used to deplete IgG plasmablasts (*9*). Our data suggests that both, depleting B cells and reducing inflammatory antibody production will be crucial as therapeutic intervention during IBD.

In summary, our data provides detailed insight into the contribution of B cells to intestinal inflammation, revealing B cell-induced cytokines and B cell-derived antibodies as harmful contributors during IBD and identifying novel therapeutic targets.

## MATERIAL and METHODS

### Mice

*Wipf1*^*-/-*^ mice (*53*) were a kind gift from Raif Geha (Boston’s Children Hospital, Boston, USA), BALB/c WT, *RAG1*^*-/-*^, C57BL/6 WT mice were bought from Charles River and bred at the Specific Opportunist Pathogen Free (SOPF) animal facility at the TranslaTUM. Igh-J^tm1Dhu^ mice (BALB/c JHT) mice were bought from Taconic. Female mice aged 6-12 weeks were used for all experiments. Littermate controls were used whenever possible. For mixed bone-marrow (BM) chimeras, 8 – 9 week old *RAG1*^*-/-*^ recipients were irradiated with 2 × 4 Gy, and injected intravenously the day after with a mixture of 80% Balb/c JHT with 20% WT or WIP KO BM (4 × 10^6^ BM cells in total). Repopulation was determined and chimeras used 8 – 10 weeks after injection. B cells were depleted from *Wipf1*^*-/-*^ mice by i.p. administration of 250μg/ml anti-CD20 (SA271G2, Biolegend) or rat IgG2b, k Isotype control (RTK4530, Biolegend) at 6 weeks and 8 weeks of age, end analysis was performed at 10 weeks of age. To induce colitis in *RAG1*^*-/-*^ recipients, 5×10^5^ sorted naïve CD4 T cells (CD19^-^CD4^+^CD25^-^CD45RB^hi^) were injected alone or in combination with MACS purified (purity more than 97%) 3×10^5^ WT or Wipf1^-/-^ B cells. Mice were monitored for weight loss and end analysis was performed 8 weeks post injection. All mouse experiments were performed in accordance with the guidelines of the Federation of European Laboratory Animal Science Association (FELASA) and followed the legal approval of the Government of Upper Bavaria (Regierung von Oberbayern).

### Human samples

Informed, written consent was obtained from all patients and controls, with prior approval of the local ethics committee of the Faculty of Medicine of the Technical University of Munich, including two-fold pseudonymization for patient tissue and blood samples (TUM; #1926/2007, #5428/12 and 2022-297-S-KH). Blood samples were obtained from IBD patients requiring diagnostic procedures or surgery. Control blood samples were obtained from healthy donors with no prior history of IBD or other autoimmune diseases. Tissue samples for histology were obtained from IBD patients undergoing bowel resections of diseased bowel segments due to fibrotic intestinal obstruction or drug-resistant disease. Control tissue samples for histology were obtained from resection margins of surgical specimens from patients undergoing bowel resection surgery for cancer. Tissue sampling was performed in accordance with the regulations of the tissue bank of the TUM and Klinikum rechts der Isar, Munich (MTBIO).

### 16s RNA sequencing

Fecal samples were collected from the cecum of non-co-housed WT and *Wipf1*^*-/-*^ littermates. Total DNA from feces was isolated using the QiaAmp® PowerFecal® Pro DNA Kit on the QIAcube Connect device (Qiagen, Germantown, MD, USA). For 16S metagenomic analysis, variable regions V3 to V4 of the bacterial 16S ribosomal RNA gene were amplified by limited cycle PCR and subjected to the Illumina 16S metagenomic sequencing library preparation workflow according to the manufacturer’s recommendations (Illumina Inc., San Diego, CA, USA). The obtained DNA libraries were quantified, normalized, and pooled. After DNA library denaturation, samples were sequenced using MiSeq v3 sequencing chemistry with paired-end 300 bp reads on the Illumina MiSeq sequencer. Bioinformatic analysis (demultiplexing, primer/adapter/barcode removal, quality trimming, DADA2 amplicon, classification and composition analysis) was performed using the Qiime2 package (*54*). Taxonomic classification was assigned by a native Bayes classifier trained on SILVA Database Rel. 132. Data visualization through 3D PCoA plots was performed using the Emperor tool. The Jaccard coefficient was calculated as a measure of similarity of the gut microbiome composition (beta diversity) between WT and *Wipf1*^*-/-*^ mice. Microbiome sequencing data were deposited in the European Nucleotide Archive (ENA) under Project accession: PRJEB55760.

### Murine primary cell isolation and culture

Murine colons were collected, fat, luminar contents carefully removed, tissue opened longitudinally and further cut into 0.5 cm pieces. Epithelial cell removal was carried out by incubating the tissue pieces in 10 mL pre-digestion medium (HBBS-Ca^2+^ and Mg^2+^ free, 10mM HEPES, 5%FCS, 1x penicillin-streptomycin (P/S) 5mM EDTA and 1mM DTT) for 30 min rolling at 37°C. Prior to enzymatic digestion the tissue pieces were washed by vortexing in 10 mL HBSS-Ca^2+^ and Mg^2+^ rich medium containing 10mM HEPES, 1x P/S, and 5%FCS. Tissue samples were minced using scissors and incubated in 3mL digestion medium (HBSS-Ca^2+^ and Mg^2+^ rich, 10mM HEPES, 1x P/S, 5%FCS, 0.5mg/ml Collagenase D, 0.5mg/ml DNase I and 1mg/ml Dispase II) for 30 min at 37°C on a thermal bloc. The cell/tissue suspension was mechanically dissociated, passed through a 100μm cell strainer, lymphocytes purified through a Percoll gradient, harvested at the 40/80% interface and washed in ice-cold B cell medium (RPMI-1640 + GlutaMax, 10mM HEPES, 10%FCS, 1x P/S and 0.05mM 2-Mercaptoethanol). MLN and spleen primary cells were harvested either by mechanical dissociation through a 70μm cell strainer or enzymatic digestion in digestion medium for 30 min at 37°C on a thermal bloc.

Splenic and mLN naïve B cells were purified using negative B cell isolation kits yielding enriched populations of 95%–98% (Miltenyi Biotec). Splenic naïve CD4 T cells were purified using negative Naïve CD4+ T cell isolation Kit yielding a cell purity of 97%-99% (Miltenyi Biotec). For plasma cell differentiation, purified B cells were labeled in PBS with 1μM CTV (Invitrogen) for 10 minutes at 37°C. Cells were maintained in complete B cell medium (RPMI supplemented with 10%FCS, 25 mM Hepes, Glutamax, penicilin streptomycin (Invitrogen) and 1‰ß-mercaptoethanol (Sigma Aldrich). CTV labeled cells at a concentration of 2×10^6^ cells per mL were stimulated in complete B cell medium supplemented with combinations of 10μg/mL of LPS from Escherichia coli O55:B5 (Sigma) or 1μg/mL CD40L (Peprotech). Proliferation and differentiation were measured after 4 days by flow cytometry. In vitro B and T cell co-culture systems were set up with 1×10^5^ purified WTor *Wipf1*^*-/-*^ B cells (Balb/c) and 2×10^5^ purified WT naïve CD4 T cells (C57BL/6). The cultures were stimulated with 5 μg/ml LPS (Sigma) in B cell medium in the presence or absence of 10 μg/ml anti-CD86 (BioLegend) or 1 μg/ml CD40L (Peprotech) for a total of 5 days.

### Bone-marrow derived macrophage (BMDM) cultures

BM was flushed from the femur and tibia of WT mice and single-cell suspensions cultured in B cell medium supplemented with 40 ng/mL murine macrophage colony-stimulating factor, M-CSF (Immunotools) for 7 days. For murine IgG stimulation, 1×10^5^ BMDMs were primed with 20ng/ml LPS-EB (InvivoGen) ON at 37 °C. 500ng/ml purified total IgG stimulation was added the next day for 4 hrs followed by an incubation of 30 min with ATP (Merck). Culture supernatants were collected for further cytokine analysis and cells lysates were prepared for qPCR analysis. For activation with co-culture supernatants, BMDMs were activated directly with 100μl co-culture medium ON at 37 °C. Cell were harvested and lysed for qPCR analysis.

### Flow cytometric analysis

Single cell suspensions were stained for viability using the Zombie Aqua fixable viability kit (1:1000 BioLegend) and fixed with 4% paraformaldehyde (PFA, VWR international). Cells were blocked with anti-mouse CD16/32 (93, 1:200, BioLegend) and CD16.2 (9E9, 1:200, BioLegend) followed by extracellular staining for 30 min with a combination of antibodies in Table S1. For intracellular cytokine stainings, cells were cultured for 4 h at 37°C in B cell medium containing 1x Cell Activation Cocktail (without Brefeldin A) (BioLegend) and 3h with 1x Protein Transporter Inhibitor Cocktail (ThermoFisher). Post Live/Dead staining, cells were fixed, permeabilized using the Intracellular Fixation and Permeabilization Buffer Set (Thermo Fisher Scientific) and stained with the appropriate combination of antibodies in Supplementary TableS1. For staining of intracellular signaling molecules, cells were fixed with 4% formaldehyde for 10 min at room temperature and permeabilized with ice-cold methanol for 30 minutes. Cells were stained with primary antibodies indicated in Table S4. Cells were acquired on LSR Fortessa cytometer and analyzed using FlowJo software (both BD Biosciences). For flow sorting, murine splenic B cells (live CD19^+^B220^+^ cells), naïve CD4 T cells (live CD19^-^CD4^+^CD25^-^CD45RB^hi^ cells) and colonic monocytes (live CD45^+^CD19^-^CD4^-^CD11b^+^ cells) were sorted on a FACSAria III (BD biosciences) cell sorter.

### Mass cytometry

3-5×10^6^ cells per sample were stained for viability as described in Cell-ID Cisplatin (Fluidigm) with a final concentration of 2.5μM Cisplatin per sample, fixed in 2% Formaldehyde solution (w/v) methanol-free (Thermos scientific) for 10 min at room temperature (RT), washed 2x with Maxpar Cell Staining Buffer and left over night (ON) at 4°C in 1mL buffer. Cells were blocked with anti-mouse CD16/32 (93, 1:200, BioLegend) and CD16.2 (9E9, 1:200, BioLegend) followed by a surface staining with the combination of the antibodies listed in Table S2 as described in Maxpar Cell Surface Staining (Fluidigm). Intracellular and intranuclear stainings were carried out as described in Maxpar Nuclear Antigen Staining with Fresh Fix (Fluidigm) followed by a Cell-ID Intercalator-Ir staing (Fluidigm) and sample prep for measurement on a CyTOF Helios with WB injector (Fluidigm). The generated FCS files were normalized and randomized using EQ beads and concatenated. Clean-up gates for elimination of no-cell signals, live cells and single cells were conducted manually using FlowJo software (BD Bioscience). To balance the influence of markers with different dynamic ranges, we performed background subtraction and channel-based normalization. To identify distinct cell populations and clusters all markers were included in a two round unbiased high-dimensional data analysis using the FlowJo plugins, FlowSOM and UMAP.

### RNA sequencing

Purified splenic B cells from WT and *Wipf1*^*-/-*^ were stimulated with 10 μg/mL LPS for 4h and 20h at 37°C in B cell medium. At time points 0h, 4h and 20h post activation 2000 cells were sorted directly into 0.2ml semi-skirted 96 well PCR plate (Peqlab) containing 10μl 1x TCL Buffer (Qiagen) and 1%(v v^-1^) β-mercaptoethanol (Sigma-Aldrich). Samples were immediately frozen at -80°C and further processed for bulk 3′-sequencing of poly(A)-RNA (RNASeq). Barcoded cDNA of each sample was generated with a Maxima RT polymerase (ThermoFisher) using oligo-dT primer containing barcodes, unique molecular identifiers (UMIs), and an adapter. 5′ ends of the cDNAs were extended by a template switch oligo (TSO) and after pooling of all samples, full-length cDNA was amplified with primers binding to the TSO-site and the adapter (*55*). cDNA was fragmented and TruSeq-Adapters ligated with the NEBNext® Ultra™ II FS DNA Library Prep Kit for Illumina® (NEB) and 3′-end-fragments were finally amplified using primers with Illumina P5 and P7 overhangs. The library was sequenced on a NextSeq 500 (Illumina) with 75 cycles for the cDNA in read1 and 16 cycles for the barcodes and UMIs in read2. Drop-Seq tools v1.12 (*56*) were used for mapping raw sequencing data to the reference genome. The resulting UMI filtered count matrices were imported into R v3.4.4. CPM (counts per million) values were calculated for the raw data and genes having a mean CPM value less than 1 were removed from the dataset.

Gene counts were normalized using DESeq2 (v1.30.1, default settings)(*57*) and annotated using AnnotationHub (v2.22.1, Shepherd MMaL. AnnotationHub: Client to access AnnotationHub resources. R package. 2021.). Murine gene sets were retrieved using msigdbr (v7.4.1, Dolgale I. msigdbr: MSigDB Gene Sets for Multiple Organisms in a Tidy Data Format. R package. 2021). Genes encoding the five complexes of the human electron transport chain were derived from the HUGO Gene Nomenclature Committee (Gene group: Mitochondrial respiratory chain complexes)(*58*) and translated to murine gene symbols using BiomaRt (v2.46.3)(*59*). Gene Set Enrichment Analysis was performed using the GSEA Java Desktop application (BROAD) and GSEA results were visualized using the EnrichmentMap app (*60*) in Cytoscape. Heatmaps were generated using Morpheus (https://software.broadinstitute.org/morpheus).

### Human colon single-cell RNA sequencing re-analysis

Annotated gene expression matrix of immune cells ((*33*), Single Cell Portal: SCP259) were loaded, processed and re-analyzed using the *scanpy* toolkit (*ver*. 1.9.1, (*61*)).

Low-quality observations, filtered by number of genes (min.: 300, max.: 2000) and percentage of mitochondrial genes (max.: 12%) and genes expressed by fewer than 3 cells were removed. Gene counts were normalized to total counts per cell (target sum: 10^4^), logarithmized and scaled (max. value: 10). To correct for batch effects, cell were integrated by unique sample using *harmonypy* (ver. 0.0.6, (*62*) and Uniform Manifold Approximation and Projection (UMAP) for dimension reduction (nearest neighbors: 50, min. distance: 0.75, neg. edge sample rate: 9) was computed with the indicated settings.

### Histology, Immunohistochemistry, Immunofluorescence

For histology, murine colons were flushed with ice-cold PBS, cut longitudinally, rolled and fixed in 4% neutral buffered formalin for 48 hrs, dehydrated and embedded in paraffin (Leica ASP 300S) according to routine methods. Blocks were cut into 2μm thick sections and stained with Hematoxylin-Eeosin (H&E), or antibodies as indicated (CD3: clone SP7, DCS CI597R0, 1:100; B220: clone RA3-6B2, BD 550286, 1:50). IHC was performed on a Leica BondRxm using a Polymer Refine detection kit with DAB as chromogen. All slides were scanned with a Leica AT2 scanning system with ×40 magnification and evaluated with ImageScope (12.4.0.7018). H&E-stained colon slides were analyzed and scored in an unbiased manner using the scoring scheme 4 (adapted from(*18*)). Colonic crypt length was determined by an average of 20 crypts per mouse gut roll using ImageJ. Human colon tissue samples were fixed in 4%(w/v) PFA (VWR International) for 24h at RT, dehydrated, embedded in paraffin, and cut into 3.5μm thick sections. Slides were deparaffinized, rehydrated, boiled in citrate buffer (10mM pH6) for antigen retrieval, washed, and incubated in 50mM NH4Cl. Slides were blocked (2% BSA and 0.3% Triton X-100), labeled with a combination of the antibodies in Table S3 and imaged using confocal microscopy. For immunofluorescence, murine colon rolls were fixed in 4%(w/v) PFA (VWR International), equilibrated in 30%(w/v) sucrose and frozen at -80 °C in Optimal Cutting Temperature (OCT) embedding medium (Thermo Fischer Scientific). 20μm sections were cut, blocked in Blocking Buffer (PBS containing 1% (v/v) FCS, 1% (v/v) mouse serum (Jackson Immunoresearch), 0.3% (v/v) Triton X-100) and stained with antibodies in Table S3. Slides were washed in PBS, mounted with Fluoromount-G (Southern Biotech) and sealed with a clear nail polish.

Colon pieces were prepared and cleared as previously described (*37*). In short, colon was harvested, the muscle layer removed and 1 cm pieces were fixed for 1 hour at 4°C; blocked; stained with antibodies as outlined in Table S3, for 72 hours at 37°C; washed; dehydrated using ascending dilutions of isopropanol (pH∼9) (Sigma-Aldrich) and cleared using undiluted ethyl cinnamate (#W243000, Sigma-Aldrich).

Imaging was performed on an inverted Leica TCS SP8 confocal microscope with white light laser and HyD photodetectors using an HC PL APO CS2 20 × /0.75 IMM objective (zoom factor of 1) (Leica Microsystems). Images were deconvolved using the LIGHTNING module in Leica Application Suite X (Leica Microsystems). Contrast adjustment for display purposes and image analysis was performed using Imaris (Bitplane) version 9.5. Binary masking of immune cells was achieved using the Surface Creation Wizard in Imaris. Statistics of objects were then exported for cell quantification as previously described (*37*).

### Multiplex assays

Quantification of murine immunoglobulin and cytokines in blood serum, mucus scrapes and culture supernatants was carried out using commercially available LEGENDplex Multiplex Assays (BioLegend), murine serum auto-antibodies were quantified using the respective commercially available Alpha Diagnostics Intl. Kits and murine and human serum CD40L was quantified using the commercially available soluble CD40L ELISA Kits (Thermo Fisher Scientific), following the manufacturer’s procedure.

### Quantitative polymerase chain reaction

Total RNA was extracted using the RNeasy Plus Mini or Micro Kit (Qiagen). Whole tissue RNA extraction was carried out using manual tissue disruption with Qiagen. RNA purity and concentration was measured using a NanoDrop spectrophotometer (Thermo Scientific) prior to cDNA synthesis using the High-Capacity cDNA Reverse Transcription Kit (Applied Biosystems). qPCRs were carried out with the Takyon No ROX SYBR 2x MasterMix blue dTTP (Eurogentec) and the primers shown in Table S5 on the Light Cycler 480II (Roche) and gene expression was determined and normalized to *Hprt* using the 2^-ΔCt^ method.

### Immunoblot

2×10^6^ purified splenic B cells were stimulated with 10 μg/ml LPS (Sigma) in B cell medium for 20 hrs at 37°C, washed and snap-frozen on dry ice. Pellets were re-suspended in lysis buffer (150 mM NaCl with DTT, protease (PMSF, TLCK, TPCK, PIN) and phosphatase inhibitors (Nava, Glycerol-2-Phospate) and proteins separated by SDS-PAGE were transferred to either polyvinylidene difluoride or nitrocellulose membranes. Membranes were blocked and incubated in with primary antibody indicated in Table S4 at 4°C over night followed by incubation with the respective horse-radish peroxidase (HRP) coupled secondary antibodies. Blots were incubated with enhanced chemiluminescent (ECL) solution (Pierce™ ECL Western Blotting Substrate; Thermo Fisher Scientific) and exposed to photosensitive films (Amersham HyperfilmTM ECL).

### Metabolic assays

Oxygen consumption was assessed using a Seahorse XFe96 metabolite analyzer (Agilent). 1×10^6^ purified splenic B cells were plated on poly-L-lysin coated Seahorse cell culture plates, incubated in Seahorse base medium + 1mM sodium pyruvate (Thermo Fisher Scientific) + 2mM L-glutamine (Thermo Fisher Scientific) + 10mM glucose (Sigma) in a volume of 50μl for 30min, 130μl medium were added in a second step and the cells were incubated for an additional 1h. Oligomycin, FCCP and rotenone+antimycin were sequentially injected during the measurement to a final concentration of 1μM each to assess different parameters of respiration. Data was analyzed using GraphPad Prism (Dotmatics).

Glucose uptake was measured by incubating 4×10^5^ cells with 50 μM 2NBDG (Cayman Chemical) in PBS (Invitrogen) for 5 or 30 min at 37°C, washed and analyzed using flow cytometry. Mitochondrial volume was determined by incubating 4×10^5^ cells with 100nM TMRE for 30 min at 37°C, washed and analyzed using flow cytometry. For confocal microscopy, B cells were settled on coverslips coated with poly-L-lysin at 37°C. TMRE dye was added, using a final concentration of 100 nM and incubated for 30 minutes at 37 °C. Cells were measured at Leica TCS SP8 confocal microscope with White Light Laser and HyD photodetectors and analyzed with Imaris for creating surfaces for volume calculation and ImageJ segmental area measurement of generated maximal intensity projections.

### Alpha-linked 2,6-sialic Acids on Total IgG

Isolation and elution of total IgG from murine serum was performed using Dynabeads Protein G for Immunoprecipitation (Invitrogen) as per manufacturer’s protocol. Eluted samples were stored at -20 °C. Total serum IgG or Alpha-linked 2,6-sialic Acids on IgG was detected on high-affinity Nunc MaxiSorp plates (Thermo Fischer Scientific) coated with 2μg/ml F(ab’)^2^ Fragment Goat anti-mouse IgG (γ-chain specific, Jackson), diluted in PBS, ON at 4°C. After blocking with Carbo-free Blocking Solution (VectorLabs) for 1h at RT, serum dilutions and standards were added to the wells and incubated for 1h at RT. Bound total IgG was detected with Peroxidase AffiniPure Goat Anti-Mouse IgG, Fcγ-fragment specific (Jackson Immuno Research). Sialic acids were detected with Biotinylated Sambucus Nigra (Elderberry) Bark Lectin (SNA) (VectorLabs) in Carbon-free Blocking Solution and subsequently Avidin-HRP, 1000x (BioLegend) was added. Substrate TMB (Thermo Fisher Scientific) solution was added and stopped with 2M H_2_SO_4_. Absorbance was measured at 450/600nm with a Tecan Spark (Tecan). The total amount of IgG in the serum was interpolated from the mouse IgG standard curve using a linear regression curve. SNA levels were normalized to the total IgG content and expressed as ratio of SNA:IgG.

## Supplementary Materials

Figs. S1 to S7

Tables S1 to S5

## Acknowledgments

We thank Julia Jellusova, Andrea Maul-Pavicic, Adam Wahida, Dieter Saur, Ari Waismann and Nadine Hövelmeyer for valuable expertise, providing reagents and critical reading of the manuscript. We thank the biological resource unit for animal husbandry (ZPF of the MRI at TUM) and the core facility for flow cytometry (Ritu Mishra and Linda Bachmann) for excellent support. We further thank all members of the Institute for Clinical Chemistry and Pathobiochemistry for scientific discussions and support.

## Author contributions

Conceptualization: SJK

Methodology: IG, JH, MG, SJK, JA, JH, OK, BB, RR, KS, CW, PN, RSG,

KJ Investigation: IG, JH, MG, SJK, RS, AB, MW, ES, MK, RÖ

Visualization: SJK, JA

Funding acquisition: SJK, RSG, JR

Project administration: SJK

Supervision: SJK

Writing – original draft: SJK

Writing – review & editing: SJK

## Data and materials availability

The RNA and the microbiome sequencing data referenced during the study are available in a public repository from the European Nucleotide Archive (ENA) under the accession code PRJEB55760 (microbiome). All the other data supporting the findings of this study are available within the article and its Supplementary Information files and from the corresponding author upon reasonable request.

**Figure S1:**
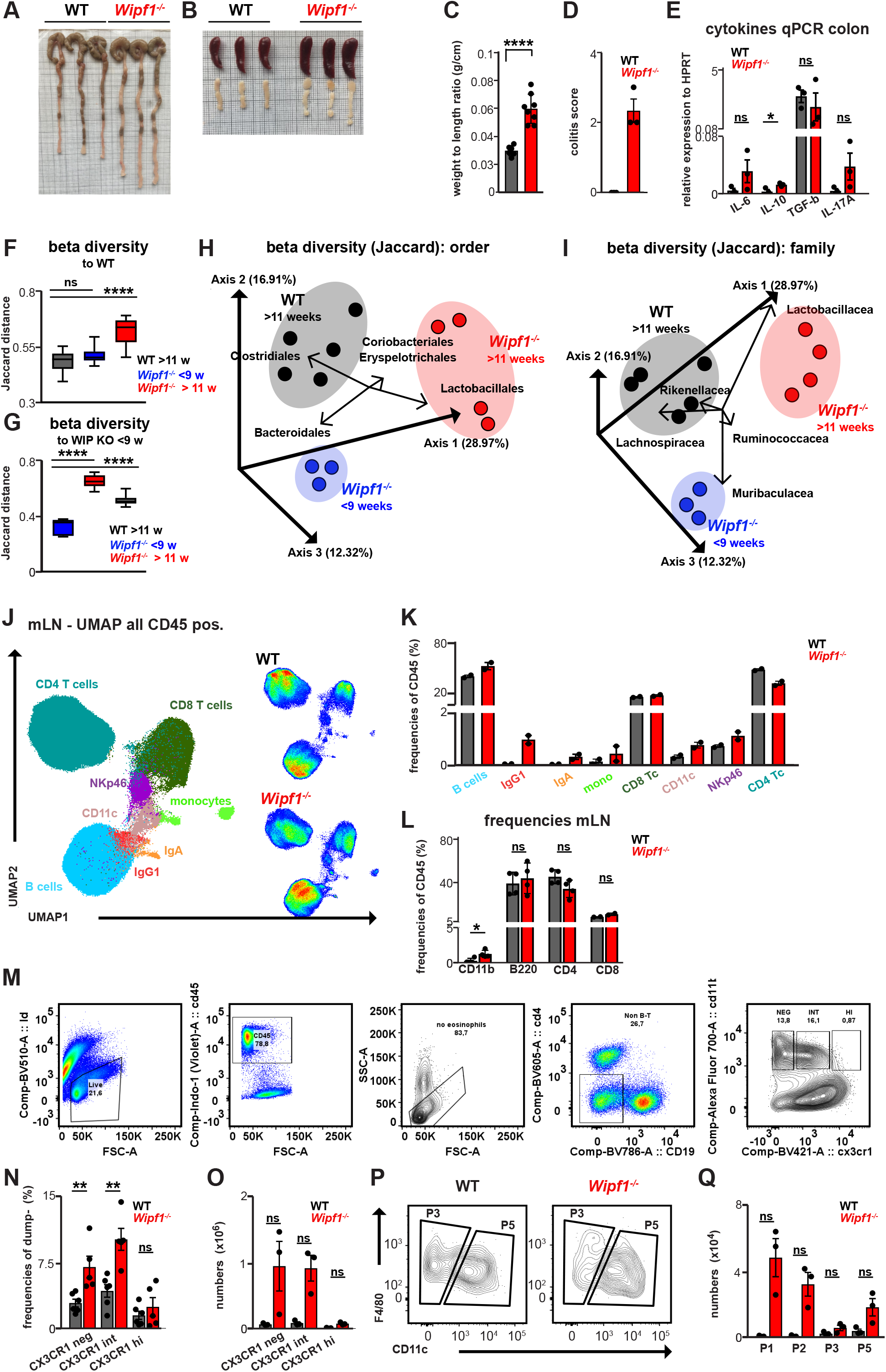
*Wipf1*^*-/-*^ mice model human chronic colonic inflammation. **(A+B)** representative pictures of (**A)** cecum and colon or (**B)** spleens and mesenteric lymph nodes isolated from 3 WT and 3 *Wipf1*^*-/-*^ mice. **(C)** Length of the proximal colon as well as the weight was measured and weight to length ratio calculated (n=8 per genotype, pooled from 4 independent experiments, mean ± SD) Statistical significance was calculated using student’s t test. (**D)** Colonic inflammation in *Wipf1*^*-/-*^ mice was scored according to (*18*). **(E)** Relative mRNA expression normalized to HPRT of indicated cytokines as determined by quantitative PCR in colonic tissue of WT and *Wipf1*^*-/-*^ mice (n=3 per genotype, pooled from 3 independent experiments, mean ± SEM) Statistical significance was calculated using student’s t test. **(F-I)** Analysis of fecal bacteria from 11 week old WT and young (<9 weeks) or older (>11 weeks) *Wipf1*^*-/-*^ mice. **(F+G)** Differences in microbial community diversities (beta diversity) between groups was analyzed using the Jaccard dissimilarity matrix (Jaccard distances). Representative data from one of two independent experiments giving similar results (Box and whisker bots (min to max)) Non-parametric ordinary one-way ANOVA with multiple comparisons was used to test for significant differences. **(H+I)** Jaccard distances (beta diversity) of **(H)** bacterial order or **(I)** bacterial family. Significant bacterial species determining the beta diversity are indicated. **(J+K)** Lymphocytes isolated from mesenteric lymph nodes of WT or *Wipf1*^*-/-*^ mice were analyzed by mass cytometry. **(J)** Uniform Manifold Approximation and Projection for Dimension Reduction (UMAP) was used to depict immune cell populations in the CD45 expressing population. FlowSOM-based immune cell populations are overlaid as color dimension **(K)** Frequencies of immune cell lineages of WT and *Wipf1*^*-/-*^ mice as defined by the FlowSOM clustering (n=2 per genotype, one representative experiment from 2 independent experiments shown). **(L)** Frequencies of immune cell lineages of WT and *Wipf1*^*-/-*^ mice as defined flow cytometry (n=4 per genotype, pooled from 3 independent experiments, mean ± SD) Statistical significance was calculated using student’s t test. **(M)** Gating strategy to determine “monocyte waterfall” subsets in single-cell suspensions isolated from the colonic lamina propria. **(N-Q)** Flow cytometric analysis of colonic “monocyte waterfall” subsets isolated from WT or *Wipf1*^*-/-*^ mice. **(N)** Frequencies and **(O)** total cell numbers of respective subsets of CD11b+ CX3CR1 expressing cells (n=5 per genotype, pooled from 2 independent experiments, or n=3 one representative experiment from 2 independent experiments shown, mean ± SEM). Statistical significance was calculated using Mann-Whitney U-test. **(P)** Representative flow blots of waterfall subsets pre-gated on CD45+, CD19-, CD4-, CD11b+, CX3CR1int., Ly6C-, MHC-II+ **(Q)** Total cell numbers of respective waterfall subsets (n=3 one representative experiment from 2 independent experiments shown, mean ± SEM). Statistical significance was calculated using Mann-Whitney U-test. *p < 0.05, **p < 0.01, ***p < 0.001, ****p < 0.0001, ns = not significant

**Fig. S2:**
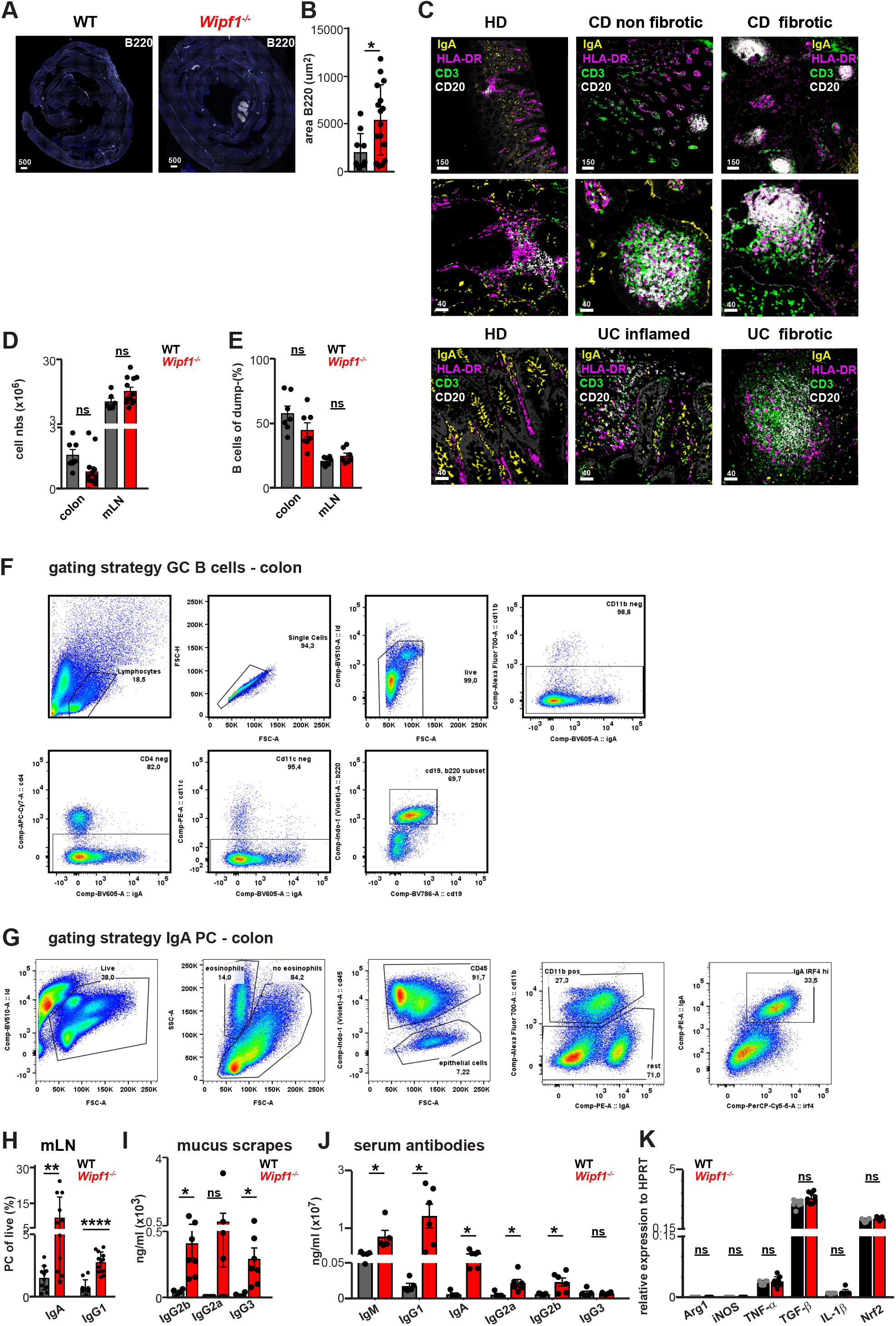
Aberrant colonic as well as systemic humoral B cell responses in *Wipf1*^*-/-*^ mice promote monocyte activation. **(A)** B220 was stained in complete colonic swiss rolls **(B)** Quantification of the total area of B220 determined using FIJI. Each dot represents one B cell follicle. Colonic swiss rolls of 4 animals per genotype were analyzed. Statistical significance was calculated using student’s t test **(C)** Immunofluorescent microscopy of representative tissue sections of colonic tissue pieces of healthy donors (HD) or non-fibrotic of fibrotic tissue of CD or inflamed and fibrotic tissue of UC patients stained for IgA plasma cells, CD3 T cells, CD20 B cells and HLA-DR (scale bars in μm). **(D)** Total cell counts from cells isolated from either colonic tissue or mesenteric lymph nodes (mLN) **(E)** Frequencies of CD19+ B cells isolated from either colonic tissue or mLN **(F+G)** Gating strategy to determine frequencies of **(F)** GC B cells or **(G)** IgA plasma cells in single-cell suspensions isolated from the colonic lamina propria. **(H)** Frequencies of indicated plasma cell populations isolated from mLN of WT or *Wipf1*^*-/-*^ mice and measured using flow cytometry. (n≥10 per genotype, mean ± SEM) Statistical significance was calculated using Mann-Whitney U-test. **(I+J)** Indicated antibody isotypes were detected in **(I)** mucus scrapes or **(J)** sera of WT or *Wipf1*^*-/-*^ mice using a multiplex assay (n≥5 per genotype, mean ± SEM) Statistical significance was calculated using student’s t test. **(K)** Relative mRNA expression of indicated mRNAs isolated from BMDMs incubated with total purified serum IgG of WT or *Wipf1*^*-/-*^ mice for 20 hrs and normalized to HPRT determined by qPCR. n≥5 sera per genotype mean ± SEM. Statistical significance was calculated using student’s t test. *p < 0.05, **p < 0.01, ***p < 0.001, ****p < 0.0001, ns = not significant

**Fig. S3:**
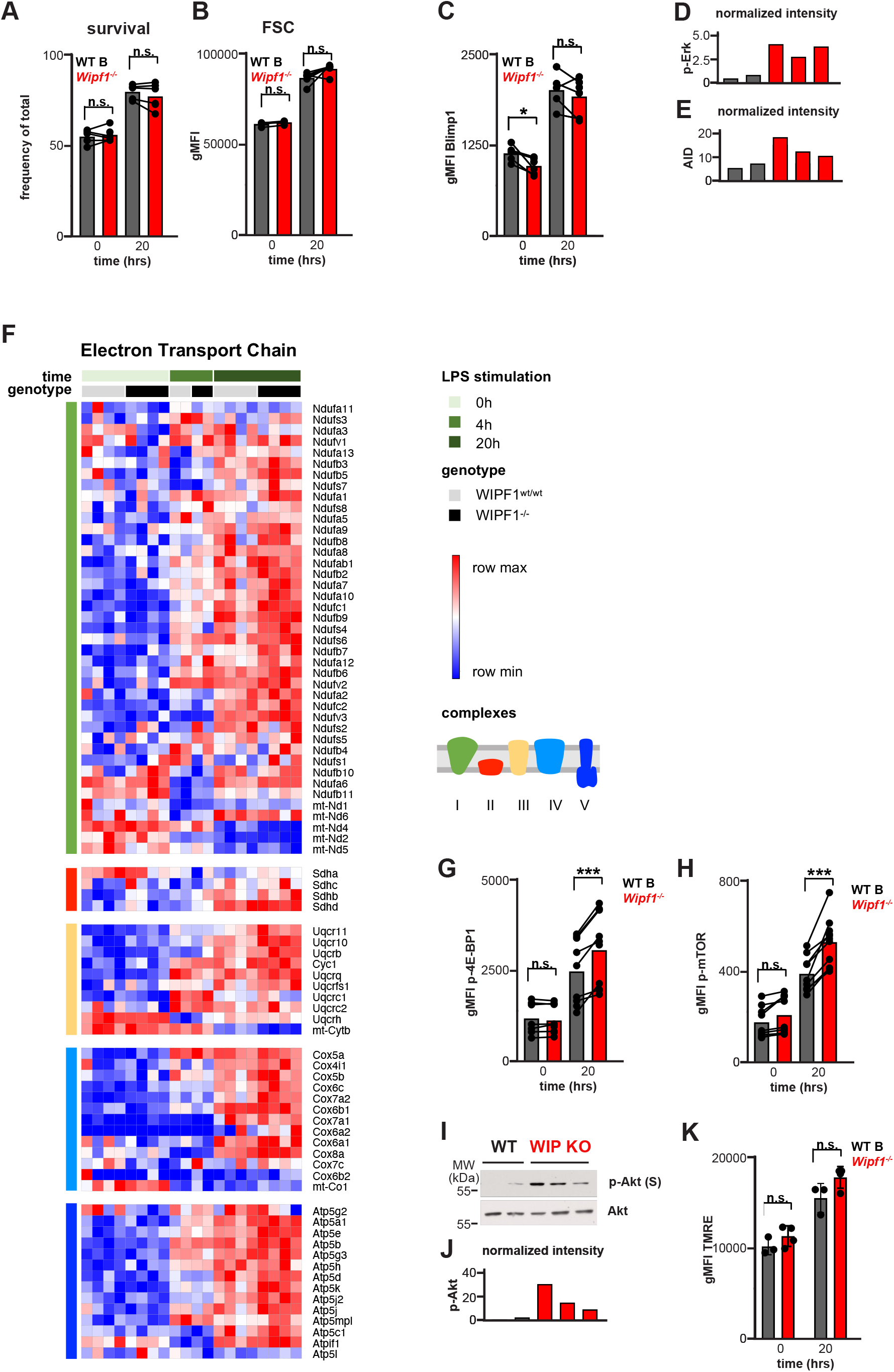
WIP deficiency metabolically primes B cells for PC differentiation. **(A-D)** Splenic WT or *Wipf1*^*-/-*^ B cells were stimulated with LPS for 20 hrs. **(A-C)** Flow cytometry analysis of **(A)** frequency of surviving cells **(B)** size of the cells **(C)** expression of Blimp1 quantified by analyzing the geometric mean fluorescent intensity (gMFI). n≥4 mice per genotype, pooled from multiple experiments. Statistical significance was calculated using paired student’s t test. **(D+E)** Quantifications of intensity of proteins normalized by densitometry to total Erk (p-Erk) or tubulin (AID). Data are representative of at least two independent experiments. **(F)** Extract of expression of genes encoding the five complexes of the electron transport chain. Color indicates z-scored DESeq2-normalized gene counts. Rows are grouped by complex and clustered hierarchically (1-pearson correlation, average linkage). n=4 mice per genotype. **(G-K)** Splenic WT or *Wipf1*^*-/-*^ B cells were stimulated with LPS for 20 hrs. **(G+H)** Quantification of the geometric mean fluorescent intensity (gMFI) analyzed by flow cytometry of phosphorylation of **(G)** 4E-BP1 and **(H)** mTORC1. n≥4 mice per genotype, pooled from multiple experiments. Statistical significance was calculated using paired student’s t test. **(I)** Immunoblot of stimulated B cells probed p-Akt as indicated. **(J)** Quantifications of intensity of proteins normalized by densitometry to total Akt. Data are representative of at least two independent experiments. **(K)** Splenic WT or *Wipf1*^*-/-*^ B cells were stimulated with LPS for 20 hrs. Flow cytometry analysis of TMRE staining quantified by analyzing the geometric mean fluorescent intensity (gMFI). n≥3 mice per genotype, pooled from multiple experiments. Statistical significance was calculated using paired student’s t test. *p < 0.05, **p < 0.01, ***p < 0.001, ****p < 0.0001, ns = not significant

**Fig. S4:**
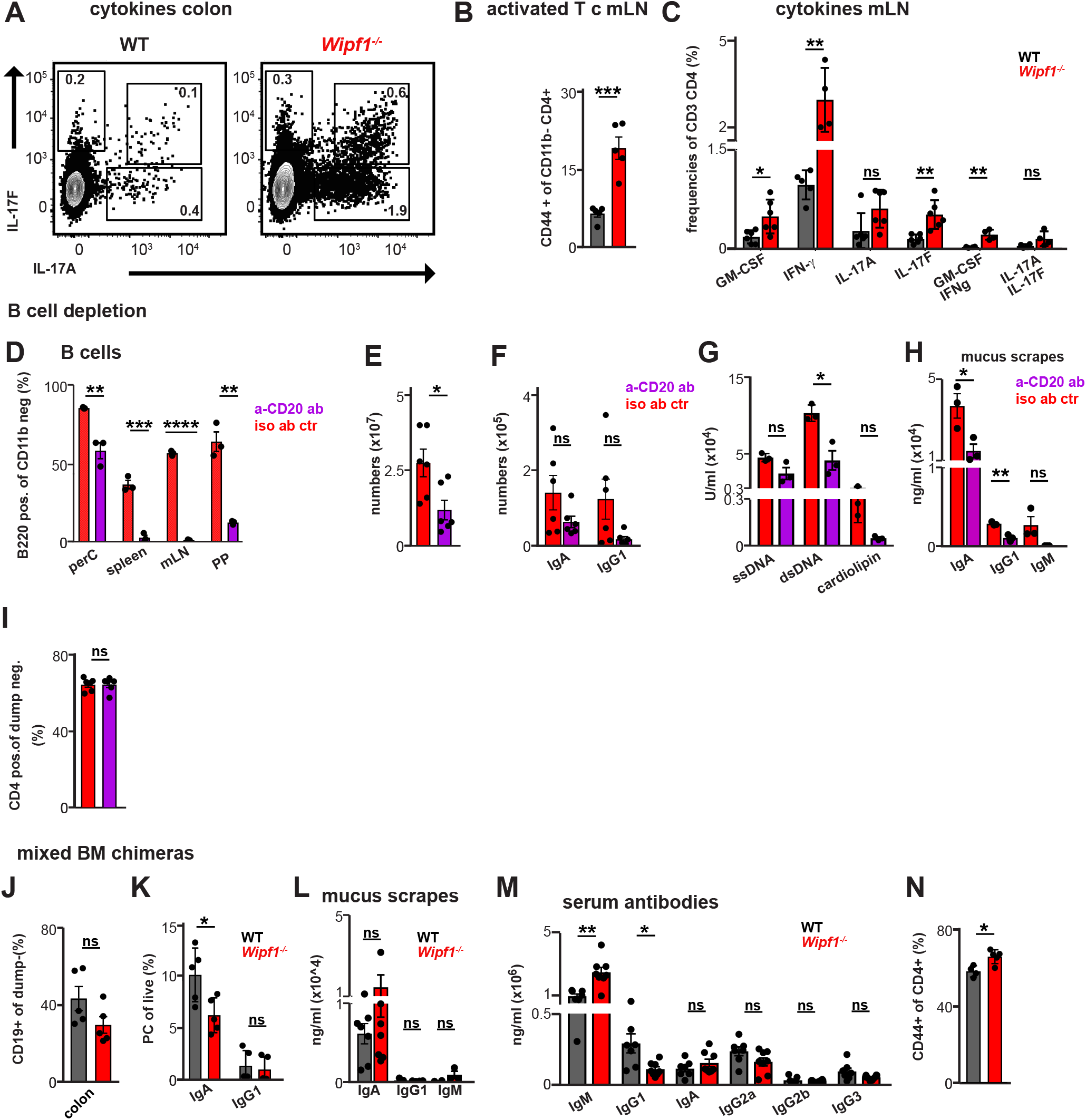
*Wipf1*^*-/-*^ B cells boost pro-inflammatory cytokine secretion by intestinal CD4 T cells. **(A)** Lymphocytes were isolated from colonic tissue of WT or *Wipf1*^*-/-*^ mice and stimulated with Brefeldin A and PMA/Ionomycin for 4 hours. Representative flow cytometric dot blots of CD4 T cells intracellularly expressing indicated cytokines. **(B)** Frequencies of CD44+ CD4 T cells of lymphocytes from mLN of WT and *Wipf1*^*-/-*^ mice. **(C)** Frequencies of intracellular staining of indicated cytokines of CD4 T cells isolated from mLN of WT and *Wipf1*^*-/-*^ mice after in vitro re-stimulation. n≥4 mice per genotype, pooled from 2 independent experiments. Statistical significance was calculated using student’s t test. **(D-I)** Lymphocytes were isolated from mLN of *Wipf1*^*-/-*^ mice treated with either isotype control antibody or anti-CD20 antibody to deplete B cells **(D)** Frequencies of B cells in indicated organs after a-CD20 antibody treatment (n=3 per genotype, data representative of 2 independent experiments) **(E)** total cell numbers isolated from mLNs. **(F)** Lymphocytes isolated from mLNs were analyzed by flow cytometry. Total cell numbers of IRF4 expressing CD45+ cell IgA and IgG1 PC subsets depicted. **(G)** Indicated serum auto-antibodies detected by ELISA. **(J)** Indicated antibody isotypes detected in mucus scrapes by a multiplex assay. **(I)** Frequencies of CD4 T cells in mLNs **(D,G+H)** n=3 per genotype, data representative of 2 independent experiments **(E,F+I)** n≥5 per genotype, pooled from 2 independent experiments. Statistical significance was calculated using student’s t test. **(J+K)** Lymphocytes were isolated from colonic tissue of JHT-WT or JHT-*Wipf1*^*-/-*^ mixed BM chimeric mice. **(J)** Frequencies of B cells. **(K)** Frequencies of PCs intracellulary expressing the indicated antibody isotypes **(L+M)** Indicated antibody isotypes detected in mucus scrapes of JHT-WT or JHT-*Wipf1*^*-/-*^ mixed BM chimeric mice by a multiplex assay **(M)** Indicated serum auto-antibodies detected by ELISA experiments **(N)** Lymphocytes were isolated from colonic tissue of JHT-WT or JHT-*Wipf1*^*-/-*^ mixed BM chimeric mice. Frequencies of CD44+ CD4 T cells are depicted. **(J,K+N)** n=5 per genotype, data representative of 2 independent experiments **(L+M)** n=7 per genotype, pooled from 3 independent experiments. Statistical significance was calculated using student’s t test. *p < 0.05, **p < 0.01, ***p < 0.001, ****p < 0.0001, ns = not significant

**Figure S5:**
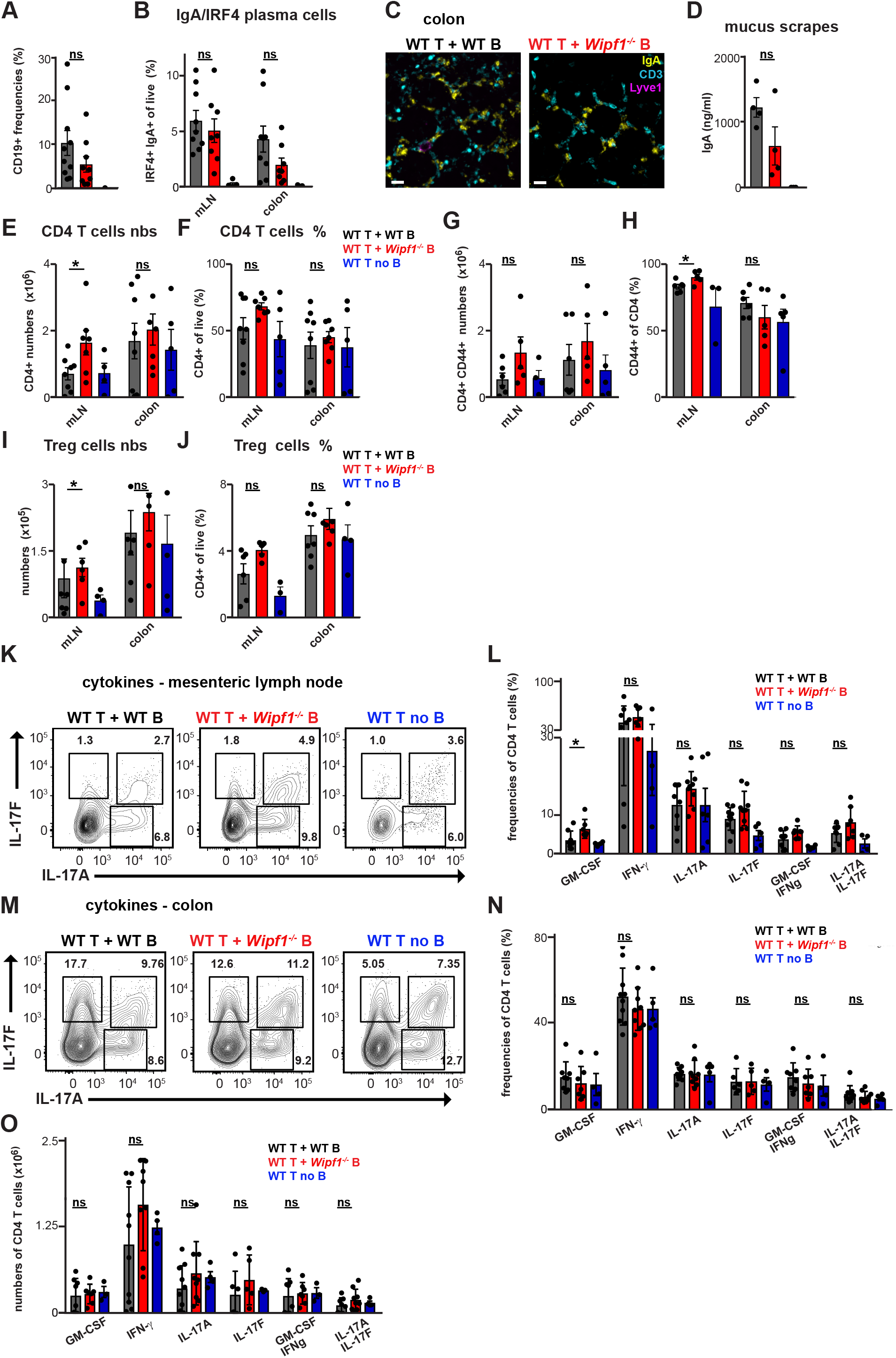
*Wipf1*^*-/-*^ B cells are sufficient to enhance pro-inflammatory cytokine production in WT CD4 T cells in an acute model of colitis. **(A)** Frequencies of B cells isolated from mLN of WT or *Wipf1*^*-/-*^ mice **(B)** Frequencies of IgA+ PC isolated from mLNs or colonic tissue. **(C)** Confocal microscopy of colonic IgA PC (yellow), CD3 T cells (cyan) and Lyve-1 positive vessels (magenta) in mice after adoptive transfer as indicated in Figure 3A. **(D)** Concentration of intestinal IgA antibodies detected in mucus scrapes by a multiplex assay. **(E)** Total numbers and **(F)** frequencies of CD4 T cells isolated from mLNs or colonic tissue. **(G)** Total numbers and **(H)** frequencies of CD44 expressing CD4 T cells isolated from mLNs or colonic tissue. **(I)** Total numbers and **(J)** frequencies of regulatory T cells (CD4+, FoxP3+) T cells isolated from mLNs or colonic tissue. **(K-O)** Lymphocytes were isolated from **(K+L)** mLN or **(M-O)** colonic tissue and stimulated with Brefeldin A and PMA/Ionomycin for 4 hours. **(K+M)** Representative flow cytometric dot blots of CD4 T cells intracellularly expressing indicated cytokines. **(L)** Frequencies of CD4 T cells expressing indicated cytokines after in vitro re-stimulation and intracellular staining. **(N)** Frequencies and **(O)** total cell numbers of CD4 T cells expressing indicated cytokines after in vitro re-stimulation and intracellular staining. n≥4 per genotype, pooled from 3 independent experiments (all mean+/-SEM). Statistical significance was calculated using student’s t test. *p < 0.05, **p < 0.01, ***p < 0.001, ****p < 0.0001, ns = not significant

**Figure S6:**
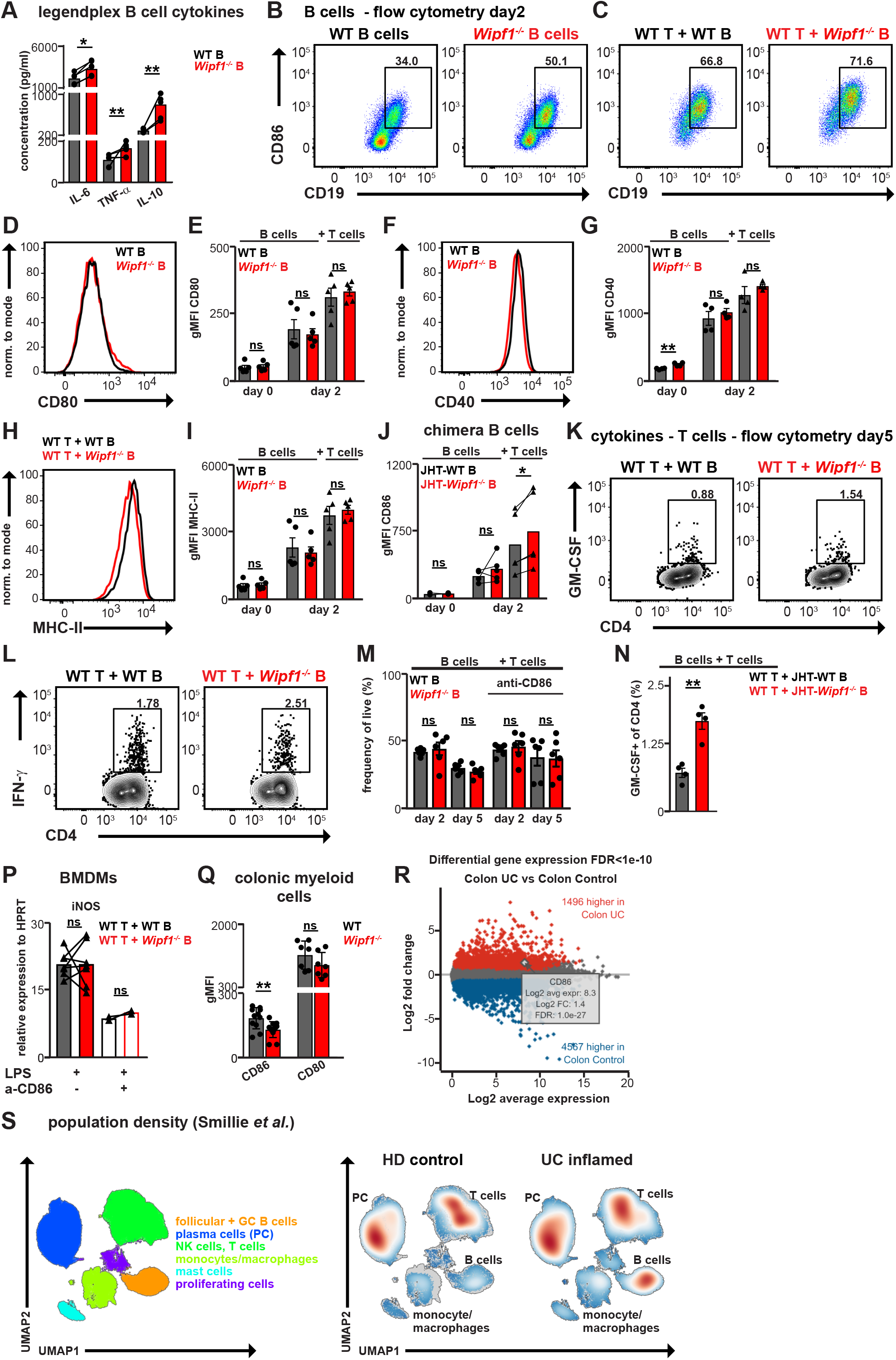
B cell mediated co-stimulation via CD86 enhances CD4 T cell cytokine production. **(A)** Purified mLN B cells were stimulated with LPS for 24 hrs. Indicated cytokines in the culture supernatants were analyzed using a multiplex assay. n≥3 mice per genotype, pooled from multiple experiments. **(B+C)** Flow cytometry analysis of CD86 expression on B cells stimulated for 48 hrs with LPS in the **(B)** presence or **(C)** absence of T cells. **(D-I)** Flow cytometry analysis of **(D+E)** CD80 **(F+G)** CD40 or **(H+I)** MHC-II expression on mLN WT or *Wipf1*^*-/-*^ B cells stimulated with LPS for 48 hrs in the presence of T cells. **(E,G,I)** Indicated surface marker expression quantified by analyzing the geometric mean fluorescent intensity (gMFI). **(D-I)** n≥4 mice per genotype, pooled from multiple experiments. **(J)** CD86 expression shown as gMFI on purified splenic B cells of JHT-WT or JHT-*Wipf1*^*-/-*^ mixed BM chimeric mice stimulated with LPS for 48 hrs in the presence or absence of T cells analyzed by flow cytometry. n≥4 mice per genotype, two independent experiments. **(K+L)** Exemplary dot blots of **(K)** GM-CSF or **(L)** IFN-γ expressing CD4 T cells stimulated with Brefeldin A and PMA/Ionomycin for 4 hours, stained and analyzed by flow cytometry at day 5 of co-culture. **(M)** Frequencies of living B cells in co-cultures at day 2 and 5 in the presence of T cells. n≥5 mice per genotype, pooled from multiple experiments. **(N)** Frequencies of GM-CSF expressing WT CD4 T cells co-cultured with B cells isolated from splenic JHT-WT or JHT-*Wipf1*^*-/-*^ mixed BM chimeric mice, stimulated with Brefeldin A and PMA/Ionomycin for 4 hours, stained and analyzed by flow cytometry at day 5 of co-culture. n=4 mice per genotype, pooled from 2 independent experiments. **(P)** BMDMs were incubated with indicated co-culture supernatants (n=5 experiments) for 20 hrs. Relative mRNA expression of *iNOS* normalized to HPRT determined by qPCR. **(Q)** CD86 and CD80 expression shown as gMFI on ex vivo isolated colonic CD11b^+^ monocytes of WT or *Wipf1*^*-/-*^ mice analyzed by flow cytometry. n≥5 mice per genotype. **(R)** Differential gene expression of CD86 from bulk RNA sequencing comparing tissue from UC patients to HD control (data from IBD TaMMA (ibd-meta-analysis.herokuapp.com)). **(S)** Single-cell RNA sequencing (scRNA-seq) analysis demonstrates an increase in B cells in colonic biopsies of inflamed UC patients compared to healthy tissue (data extracted from *33*). Statistical significance was calculated using **(A, J, P)** paired student’s t test or **(E, G, I, M, N, Q)** student’s t test. *p < 0.05, **p < 0.01, ***p < 0.001, ****p < 0.0001, ns = not significant

**Table S1:** antibodies for flow cytometry.

| Target species | Antigen | Fluorophore | Clone | Company |
| --- | --- | --- | --- | --- |
| mouse | B220/CD45R | BV421 | RA3-6B2 | BioLegend |
| mouse | B220/CD45R | BUV395 | RA3-6B2 | BD Biosceinces |
| mouse | Blimp-1 | AF647 | 5E7 | BioLegend |
| mouse | CCR9 | FITC | CW-1.2 | BioLegend |
| human/ mouse | CD11b | AF700 | M1/70 | BioLegend |
| mouse | CD11c | PE/Biotin | N418 | BioLegend |
| mouse | CD138 | PerCP-Cy5.5 | 281-2 | BioLegend |
| mouse | CD19 | BV785 | 6D5 | BioLegend |
| mouse | CD25 | PE | PC61.5 | eBioscience |
| mouse | CD3e | APC-Cy7/BV421 | 145-2C11 | BioLegend |
| mouse | CD4 | APC-Cy7<br>/BV605/FITC/BV785/B<br>V421 | GK1.5 | BioLegend |
| mouse | CD40 | FITC | HM40-3 | BioLegend |
| human/ mouse | CD44 | APC-Cy7 | IM7 | BioLegend |
| mouse | CD45 | BUV395 | 30-F11 | BD Biosceinces |
| mouse | CD45R/B220 | BUV395 | RA3-6B2 | BD Biosceinces |
| human/mouse | CD45R/B220 | BV785 | RA3-6B2 | BioLegend |
| mouse | CD45RB | APC | c363-16a | BioLegend |
| mouse | CD62L | APC | MEL-14 | eBioscience |
| mouse | CD64 | PerCP-Cy5.5 | X54-5/7.1 | BioLegend |
| mouse | CD80 | PE/APC | 16-10A1 | BioLegend |
| mouse | CD86 | PE/PerCP-Cy5.5 | GL-1 | BioLegend |
| mouse | CD8a | APC/PerCP-Cy5.5/BV421 | 53-6,7 | BioLegend |
| mouse | CX3CR1 | BV421 | SA011F11 | BioLegend |
| mouse | F4/80 | PE/Cy7 | BM8 | eBioscience |
| mouse | F4/80 | BV 421 | BM8 | BioLegend |
| mouse | Fas/CD95 | PE-Cy7 | Jo2 | BD Biosceinces |
| mouse | FoxP3 | PE | FJK-16a | eBioscience |
| human/mouse | GL7 | AF 488 | GL7 | BioLegend |
| mouse | GM-CSF | PerCP-Cy5.5/PE | MP1-22E9 | BioLegend |
| mouse | IFN-γ | BV421/FITC/APC/Cy7 | XMG1.2 | BioLegend |
| mouse | IgA | APC | 11-44-2 | Southern Biotech |
| mouse | IgA | AF555/AF488 | Polyclonal | Southern Biotech |
| mouse | IgA | PE/FITC | 11-44-2 | Southern Biotech |
| mouse | IgA | Biotin | RMA-1 | BioLegend |
| mouse | IgG1 | APC/FITC | A85-1 | BD Biosceinces |
| mouse | IgM | eF 450 | II/41 | BioLegend |
| mouse | IL-1b pro-form | PE | NJTEN3 | eBioscience |
| mouse | IL-10 | APC/FITC | JES5-16E3 | eBioscience |
| mouse | IL-17A | eF660/PE | eBio17B7 | eBioscience |
| mouse | IL-17F | AF647/PE/AF488 | 9D3.1C8 | BioLegend |
| mouse | IL-6 | PerCP-Cy5.5/APC | MP5-20F3 | BioLegend |
| mouse | IRF4 | eF660 | 3E4 | eBioscience |
| mouse | IRF4 | PerCP-Cy5.5 | 3E4 | BioLegend |
| mouse | Ly-6C | FITC | HK1.4 | BioLegend |
| mouse | MHC II | FITC | M5/114.15.2 | BioLegend |
| mouse | MHCII | APC-Cy7 | M5/144.15.2 | eBioscience |
| mouse | NKp46 | PerCP-Cy5.5 | 29a1.4 | BioLegend |
| mouse | p4EBP1 (Thr 36, Thr45) | eF 660 | V3NTY24 | eBioscience |
| human/ mouse | pAKT (S473) | eF450 | SDRNR | eBioscience |
| human/ mouse | pmTOR | eF450 | MRRBY | eBioscience |
| human/mouse | p38 |  | rabbit | CellSignalling |
| human/mouse | p-p38 MAPK |  | rabbit | CellSignalling |
| mouse | RORyt | APC | b2d | eBioscience |
| mouse | T-bet | PE | eBio4B10 | eBioscience |
| mouse | TACI/CD267 | PE | 8F10 | BioLegend |
| mouse | TNF-alpha | PerCP-Cy5.5/AF488/APC/Cy7 | MP6-XT22 | BioLegend |
|  | TMRE | Emission 576nm |  | Thermofischer Scientific |

**Table S2:**
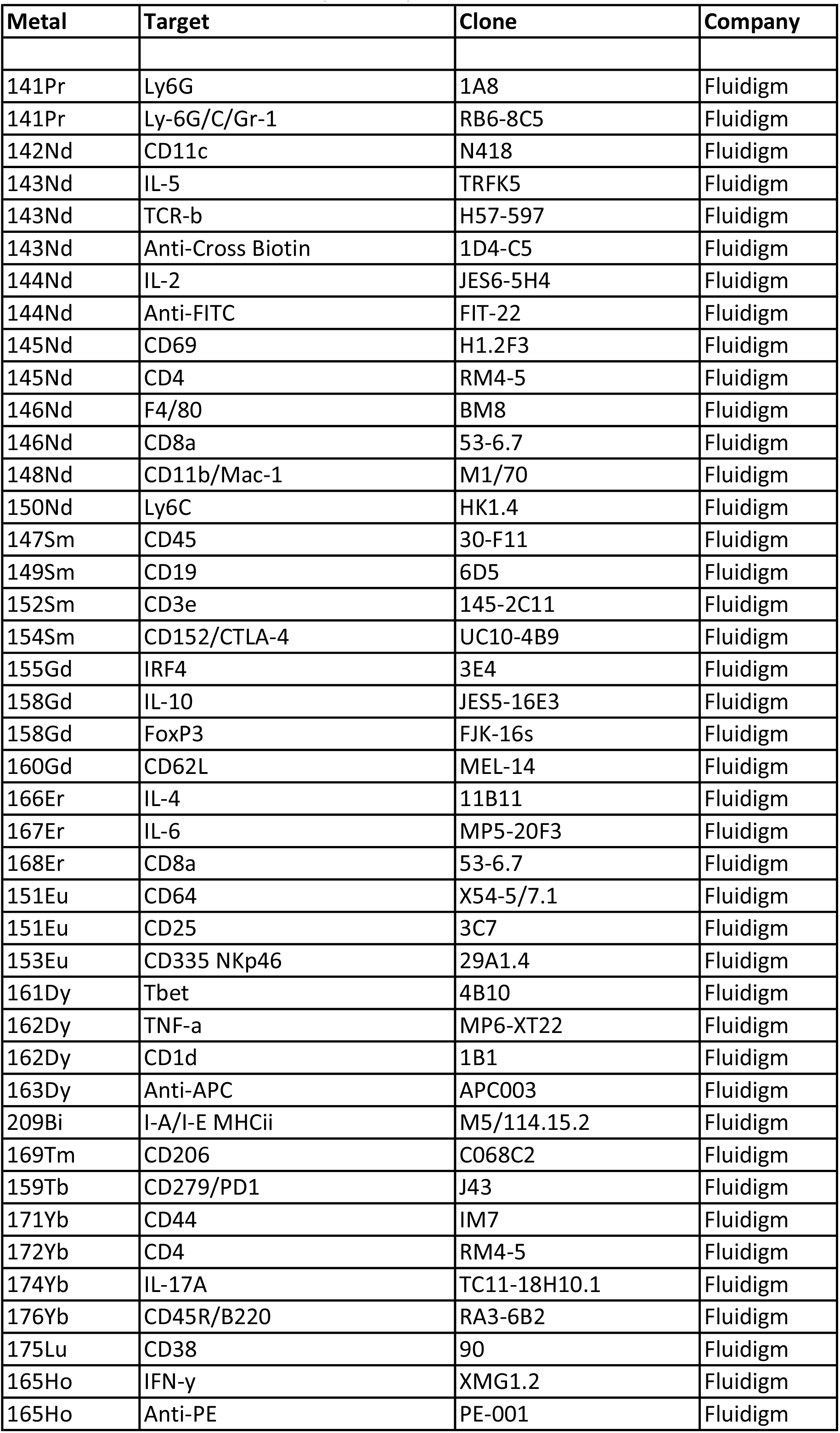
antibodies for mass cytometry.

**Table S3:** antibodies for confocal microscopy.

| Target species | Antigen | Fluorophore | Clone | Company |
| --- | --- | --- | --- | --- |
| mouse | MHCii | BV421/AF488 | M5/114.15.2 | BioLegend |
| mouse | CD11b | AF488/AF700 | M1/70 | BioLegend |
| mouse | IgA | AF555 | polyclonal | Southern Biotech |
| mouse | CD3 | AF594 | 17A2 | BioLegend |
| mouse | Lyve-1 | AF488/eF660 | ALY7 | BioLegend |
| human | IgA | AF555 | polyclonal | Southern Biotech |
| human | HLA-DR | BV421 | L243 | BioLegend |
| human | CD3 | AF594 | UCHT1 | BioLegend |
| human | CD20 | AF488 | L26 | BioLegend |

**Table S4:** antibodies for immunoblot/phospho-flow.

| Target species | Antigen | Source species | Product number | Company |
| --- | --- | --- | --- | --- |
| mouse | p-Erk p-44/p42 MAPK (T202/Y204) | rabbit | #9101 | Cell Signaling |
| mouse | Erk1/2 p42/44 MAPK | rabbit | #9102 | Cell Signaling |
| mouse | AID (L7E7) | mouse | #4975 | Cell Signaling |
| mouse | a/b tubulin | rabbit | #2148 | Cell Signaling |
| mouse | p-Akt (S473) | rabbit | #4058 | Cell Signaling |
| mouse | Akt | rabbit | #9272 | Cell Signaling |
| mouse | p-p38 | rabbit | #4511 | Cell Signaling |
| mouse | p-mTORC1 | rabbit | 48-9718-42 | Invitrogen |
| mouse | p-4E-BP1 | rabbit | #5123 | Cell Signaling |
| mouse | Blimp1 | rabbit | #9115 | Cell Signaling |

**Table S5:**
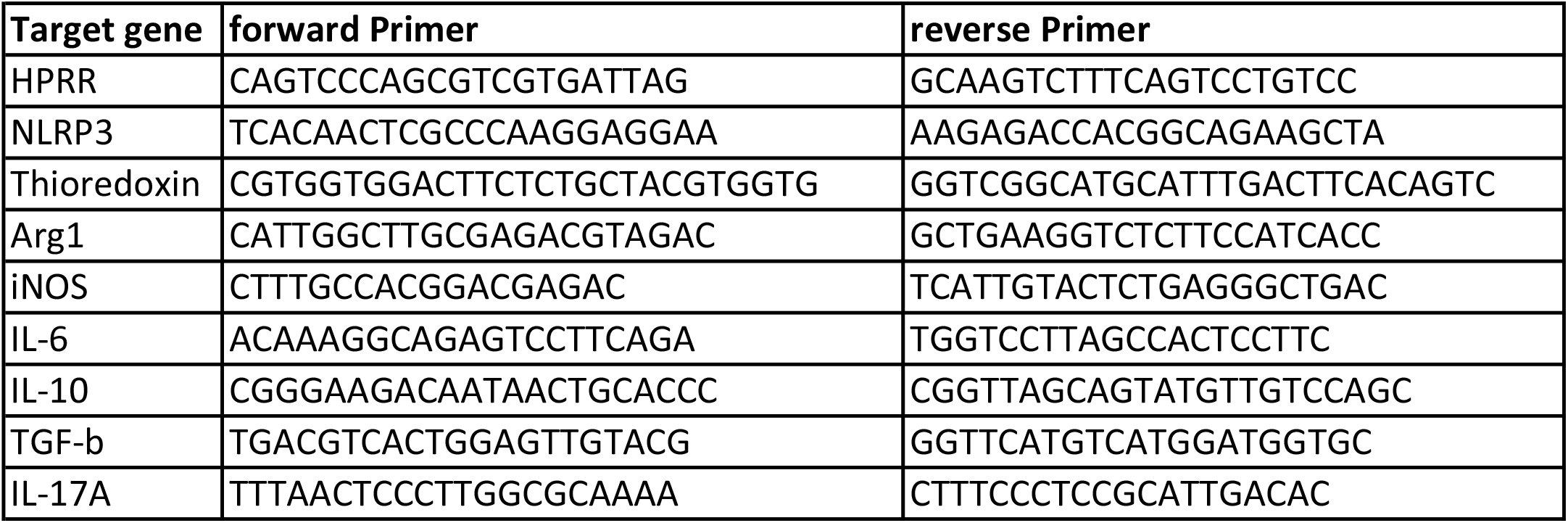
qPCR primers.

